# Nucleoid associated proteins and their effect on *E. coli* chromosome

**DOI:** 10.1101/2020.11.05.369934

**Authors:** Ankit Gupta, Abdul Wasim, Jagannath Mondal

## Abstract

A seemingly random and disorganized bacterial chromosome, in reality, is a well organized nucleus-like structure, called the nucleoid, which is maintained by several nucleoid associated proteins(NAPs). Here we present an application of a previously developed Hi-C based computational method to study the effects of some of these proteins on the *E. coli* chromosome. Simulations with encoded Hi-C data for mutant, hupAB deficient, *E. coli* cells, revealed a decondensed, axially expanded chromosome with enhanced short range and diminished long range interactions. Simulations for mutant cells deficient in FIS protein revealed that the effects are similar to that of the hupAB mutant, but the absence of FIS led to a greater disruption in chromosome organization. Absence of another NAP, MatP, known to mediate Ter macrodomain isolation, led to enhanced contacts between Ter and its flanking macrodomains but lacked any change in matS sites’ localization. Deficiency of MukBEF, the only SMC complex present in *E. coli*, led to disorganization of macrodomains. Upon further analysis, it was observed that the above mutations do not significantly impact the local chromosome organization (~ 100 Kb) but only affect the chromosome on a larger scale (>100 Kb). These observations shed more light on the sparsely explored effects of NAPs on the overall chromosome organization and helps us understand the myriad complex interactions NAPs have with the chromosome.

## Introduction

For decades, the structure and packing of chromosomes inside small cells and nucleus remained a mystery. Due to large size of eukaryotic cells, the mechanism of DNA folding/packing was first understood in eukaryotes. In bacteria, it was believed that their genome is largely unstructured due to lack of DNA structuring factors such as histone proteins as in case of eukaryotes[1]. But at the end of 20th century evidence started accumulating suggesting the presence of proteins involved in nucleus like structure management in bacteria[2]. The common eubacteria *E. coli*, has single circular chromosome with genomic size of ~ 4.64 Mbps. Although the contour length is ~ 1.5mm, it is packed compactly into a nucleus like structure called as nucleoid, inside a 1.5 − 2.5*μ*m long cell.

Initial studies to determine the nucleoid structure in bacteria include light and electron microscopic techniques. These investigations showed that, in gently lysed bacteria, the nucleoid is folded into a ‘rosette’ like model with a central dense core and several loops coming out of it [3, 4, 5]. And to obtain the fully relaxed DNA, a few tens of nicks were required[6]. Biochemical studies, with *E. coli*, to determine the nucleoid composition suggest that the whole nucleoid is composed of ~ 80% DNA, ~ 10% RNA and, ~ 10% protein[7, 6]. Altogether, these initial experiments provided the first insights into the structure of nucleoid, which is, in condensed form, having rosette like structure and arranged into topologically independent loops and sustained with the help of proteins and RNA molecules.

Previously, the protein component of the nucleoid was believed to be majorly RNA polymerase [6]. But later studies found more than 200 potential DNA binding proteins in *E. coli* cells[8]. Out of those potential proteins, 10-20 were suggested to be responsible for nucleoid organization[9]. These proteins were called nucleoid associated proteins (NAPs). Through DNA binding study, it was found that many of the NAPs had specific binding sites whereas others can bind non-specifically throughout the chromosome[9]. However, those non-specific binding proteins had a preference for AT rich segments. In an another experiment, using immunofluorescence microscopy, distribution of several of those NAPs was determined[10]. Many NAPs showed uniform distribution throughout the chromosome and others showed locus specific binding. NAPs which had specific binding sites showed the irregular distribution whereas non-specifically binding NAPs were uniformly distributed in the cells.

HU (Heat Unstable nucleoid protein) is one such protein that binds non-specifically to the whole bacterial chromosome and shows uniform distribution. It plays an important role in DNA compaction and negative supercoiling[11] of the chromosome. FIS (Factor for Inversion Stimulation) is another important nucleoid associated protein. It binds to DNA in both, specific and non-specific manner, and plays a role in gene regulation as well as nucleoid organization[12, 13]. Further investigations led to the discovery of macrodomains inside the *E. coli* chromosome[14, 15, 16]. Currently, it is believed that there are macrodomain specific proteins which govern their condensation, and isolation from other macrodomains. In search of such proteins, Macrodomain Ter Protein (MatP) was discovered[17] which has 23 specific binding sites in the Ter macrodomain (called as matS)[17]. *E. coli* also has an SMC (Structure Maintenance of Chromosome) complex called as MukBEF which is similar to condensin proteins in eukaryotes. MukBEF was first identified as a protein crucial for chromosome segregation in *E. coli* cells and required for correct positioning of Ori macrodomain[18, 19, 20]. Also, it is known that MukBEF complex facilitate DNA loops formation and helps in its compaction[21].

In current work, we have used a recently introduced simulation method, to determine the *E. coli* chromosome architecture[22], and attempted to elucidate the role of these NAPs on overall structure of the *E. coli* nucleoid. Here, we have made use of the recently published Hi-C data for *E. coli* NAPs mutants[23] to quantify the structural changes in the chromosome ensemble.

## Materials and Methods

To generate a model for the *E. coli* chromosome, we have used a previously reported protocol[22]. As per the protocol, we generated 928 bead polymers, with volume exclusions, where each bead corresponds to 5 Kb which is also the resolution of the Hi-C contact probability matrix. Then we converted Hi-C data[23] to a distance matrix using Eqn. (1), where *P* is the Hi-C contact probability matrix and *D* is the distance matrix. We used the distances from *D* to incorporate harmonic restraints into the polymer using Eqn. (2).

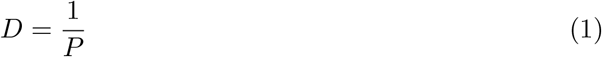

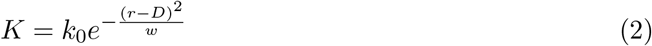

In Eqn. (2), *k*_0_ is a parameter that decides the maximum strength of a Hi-C bond. It does not affect the organization of the polymer in any way. *k*_0_ = 10 has been used for our simulations, which is an arbitrary choice. *k*_0_ is generally kept much smaller than the bonds connecting adjacent beads of the polymer model. *w* is another parameter that decides how steep the gaussian should fall. It needs to be optimized to generate a simulated contact probability matrix that has the highest correlation with the experimental one (see SI Table S1).

Figure 1 shows the overall scheme of the whole procedure to develop chromosome models for wildtype as well as mutant *E. coli*, further details regarding the methods can be found elsewhere[22].

**Figure 1:**
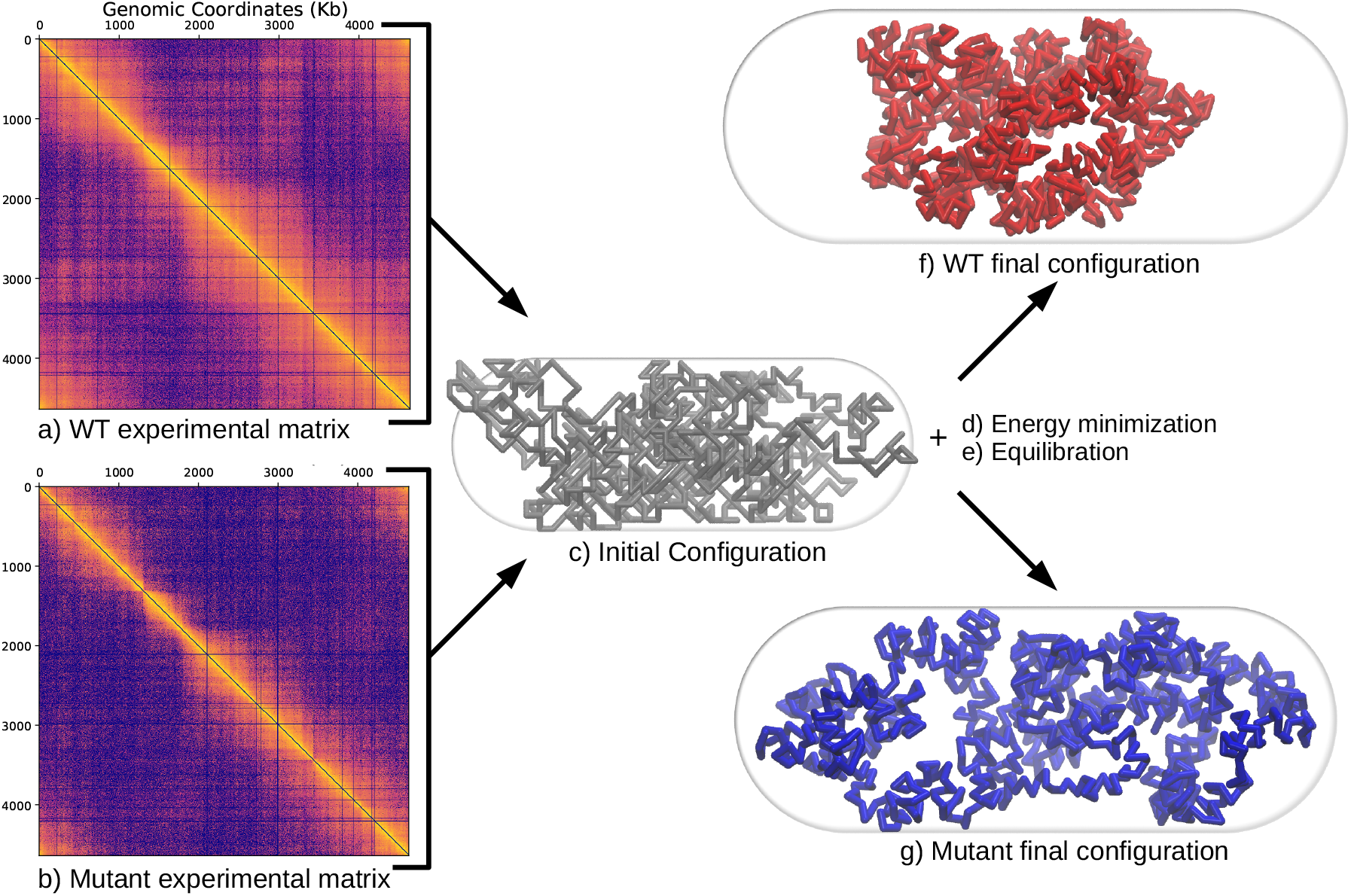
A schematic of the method used to generate an ensemble of final chromosome structures. **a)** Contact probability matrix from Hi-C experiment for wildtype (WT) *E. coli*. **b)** Contact probability matrix from Hi-C experiment for mutant *E. coli*. **c)** Initial configurations generated along with the **d)** energy minimization step and **e)** equilibration. **f,g)** Chromosome after equilibration of an energy minimized conformation for WT and mutant, respectively. Average bond distances from equilibrium conformations were converted to a contact probability matrix and the simulated and the experimental matrices are compared and the simulated matrix is used for further analysis.

Since no quantitative information on cell sizes for mutants are known, we used the same cell dimensions, as shown in Figure 1, as it has been used for simulating the chromosome for WT cells grown in LB media at 37°C (WT37LB)[24]. This also provides us with a better way to compare the results. We performed simulations for cells where anyone of HU, FIS, MatP, and MukBEF were absent. Apart from the simulations on mutants, we also performed simulations for WT cells grown at 22°C in Minimal Media (WT22MM) and at 30°C in Minimal media (WT30MM) and WT37LB. Using the WT simulations, we attempted to quantitatively identify the extent of changes in the chromosome architecture brought upon due to one or more protein mutants. For the purpose of better contrast, correlation matrices are used throughout the results section for the WT and mutant instead of contact probability matrices but simulated contact probability matrices are used for the analysis.

For visualization and rendering of chromosome structures VMD was used[25, 26]. To calculate radius of gyration and axial/linear density (R_*g*_) a python package MDAnalysis was used[27, 28]. Asphericity was calculated as mentioned elsewhere[29]. All graphs are plotted by using Matplotlib[30].

## Results

We follow a protocol[22] used to elucidate the chromosome architecture in *E. coli* cells using the reported Hi-C data[23]. We generate 200 random polymer conformations, each having 928 beads which is equal to the dimension of the Hi-C contact probability matrix. In each such conformation, adjacent beads are connected by strong harmonic bonds (see *SI Methods*). Distance restraints are introduced using harmonic bonds between beads, whose equilibrium bond distances and force constants are calculated using Eqns. (1) and (2). We performed stochastic dynamic simulations with excluded volume interactions to generate an ensemble of polymer conformations. To validate the simulations, we generated simulated average contact probability matrices and compared it with the experimental ones.

In our previous study[22], we found that our method is capable of predicting multiple experimental results and several properties related to *E. coli* chromosome. We were able to capture the self organization of bacterial circular chromosome into different macrodomains and non-structured regions. Various genomic loci showed linear organization along long axis of the cell. Our results, for wildtype sample, showed good correlation with experimental results, such as recombination frequencies and interfocal loci distances measured through FISH. In most of the bacteria, whole chromosome conformation is a result of complex interactions between regulatory and structural elements such as promoter and repressors, and nucleoid associated proteins (NAPs) respectively. Intrigued by the structural properties of the *E. coli* chromosome and the insights that our method can provide, in current work, we sought to determine the role of those multiple NAPs in maintaining that nucleoid architecture.

### HU Protein

HU is an important nucleoid associated protein for most of the bacteria to stabilize negative supercoiling and concomitantly nucleoid structure. It is similar to the eukaryotic histone proteins in sequence and structure[31]. HU is one of the most abundant and highly conserved NAPs in bacteria. It was found to be uniformly distributed throughout whole *E. coli* chromosome[10]. HU consists of 2 subunits *α* and/or *β* and present as both homodimer and heterodimer (hup*α*2 and hup*αβ*, respectively) in *E. coli* cells[32]. To ascertain the effect of HU mutation on whole chromosome structure we have carried out simulations by encoding Hi-C contact probability matrix of ΔhupAB mutant (*E. coli* MG1655 ΔhupAB at 37°*C* in LB media) into a previously developed method, further details of processed data are given in Table S1. Experimental and simulated Hi-C contact probability matrices of the ΔhupAB mutant (Figure S1a and S1b, respectively) are in excellent agreement with a correlation coefficient of 0.95. On comparison with respective wildtype (WT, *E. coli* MG1655 grown at 37°*C* in LB media), as shown in the Figure 2a, the heatmaps for the WT (top) and ΔhupAB mutant (bottom) show similarity at shorter length scale and diminished long range contacts in the mutant, similar to experimental Hi-C results[23].

**Figure 2:**
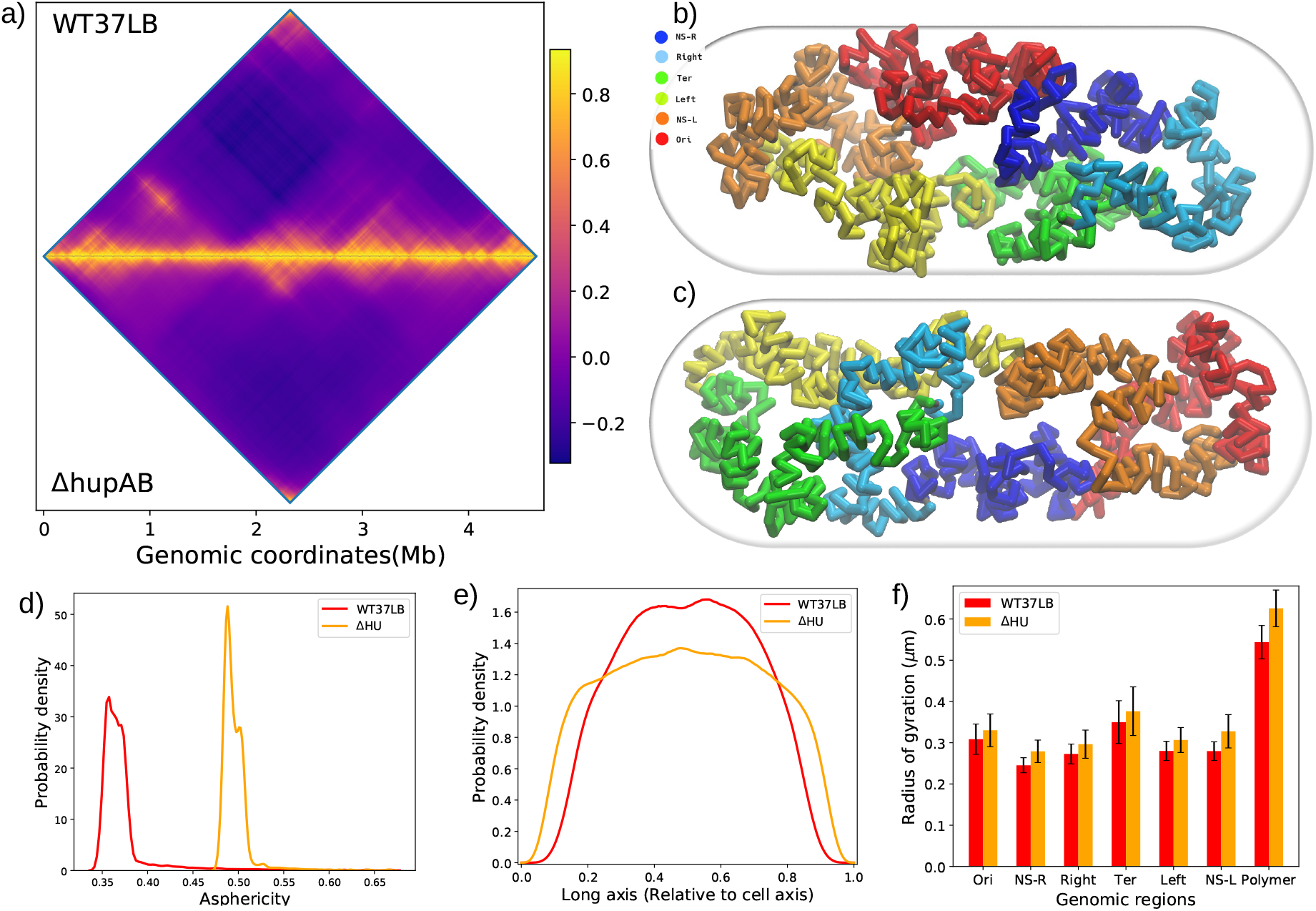
**a)** Correlation matrix of simulated WT (upper triangle) and ΔhupAB (lower triangle) contact probability matrix, **b)** Snapshot from equilibrated structure for WT *E. coli* (grown in LB at 37°C) chromosome simulation. **c)** Snapshot from equilibrated structure for ΔhupAB mutant chromosome simulation. **d)** Distribution of asphericity of WT and ΔhupAB mutant cells, **e)** Linear DNA probability density for WT and ΔhupAB mutant cells relative to cell length along the long axis (Z- axis). **f)** Radius of gyration (R_*g*_) comparison between WT and ΔhupAB mutant at same growth conditions.

Structurally, when compared with the WT (Figure 2b), whole chromosome seems to get decondensed in ΔhupAB mutant (Figure 2c) and gets dispersed across cell’s axial dimension, consistent with previous experimental observations[33]. Chromosome expansion is also reflected in the relative increase in the asphericity and axial density near the pole region in ΔhupAB mutant when compared to WT chromosome (Figure 2d and e, Table S2). Apart from overall structural changes, we also used a previously devised method R_*g*_ map [22], to see the changes in local structure of the chromosome (i.e. at 100 Kb scale). As shown in Figure S2 a and b, the changes are negligible in the mutant and they in good agreement with a correlation coefficient of 0.80. For larger genomic distances too (till 500 Kb) we observe good correlation (Table S5) which indicates although the overall structure has changed but local packaging is still similar to WT.

To get better insights on overall structure, we also calculated radius of gyration of whole chromosome and all macrodomains for ΔhupAB mutant and WT, given in Figure 2f. We find that, for each macrodomain, the R_*g*_ has increased on an average by 0.030*μ*m and for whole chromosome by 0.082*μ*m. Effect of hupAB mutation on each individual macrodomain is more apparent in their respective spatial distance versus genetic distance plot (Figure S3). Our simulations indicate that Ter macrodomain is least affected by the hupAB mutation. However, for regions in other macrodomains, especially regions separated by large genomic distances, their spatial distances increase significantly in hupAB mutant, compared to that in WT cell.

In the Hi-C experiment[23], it was found that the long range contacts (> 280 kb) were diminished and short ones (< 280 kb) were enhanced in ΔhupAB mutant, except in Ter macrodomain, where the effects were opposite[23]. Therefore, for further investigation into the changes in contact probabilities, we have calculated the difference heatmap between simulated ΔhupAB mutant and WT matrix (Figure 3a), where we have simply subtracted the WT matrix from ΔhupAB mutant matrix. In the difference heatmap, the positive/red and negative/blue contacts, therefore, imply higher contact probability in the ΔhupAB mutant and WT matrix, respectively. From the difference heatmap for ΔhupAB (Figure 3a), it is clear that the WT matrix has higher contact probability for long range contacts indicated by the off-diagonal blue regions, with few areas of enhanced (i.e. red/positive) contacts.

**Figure 3:**
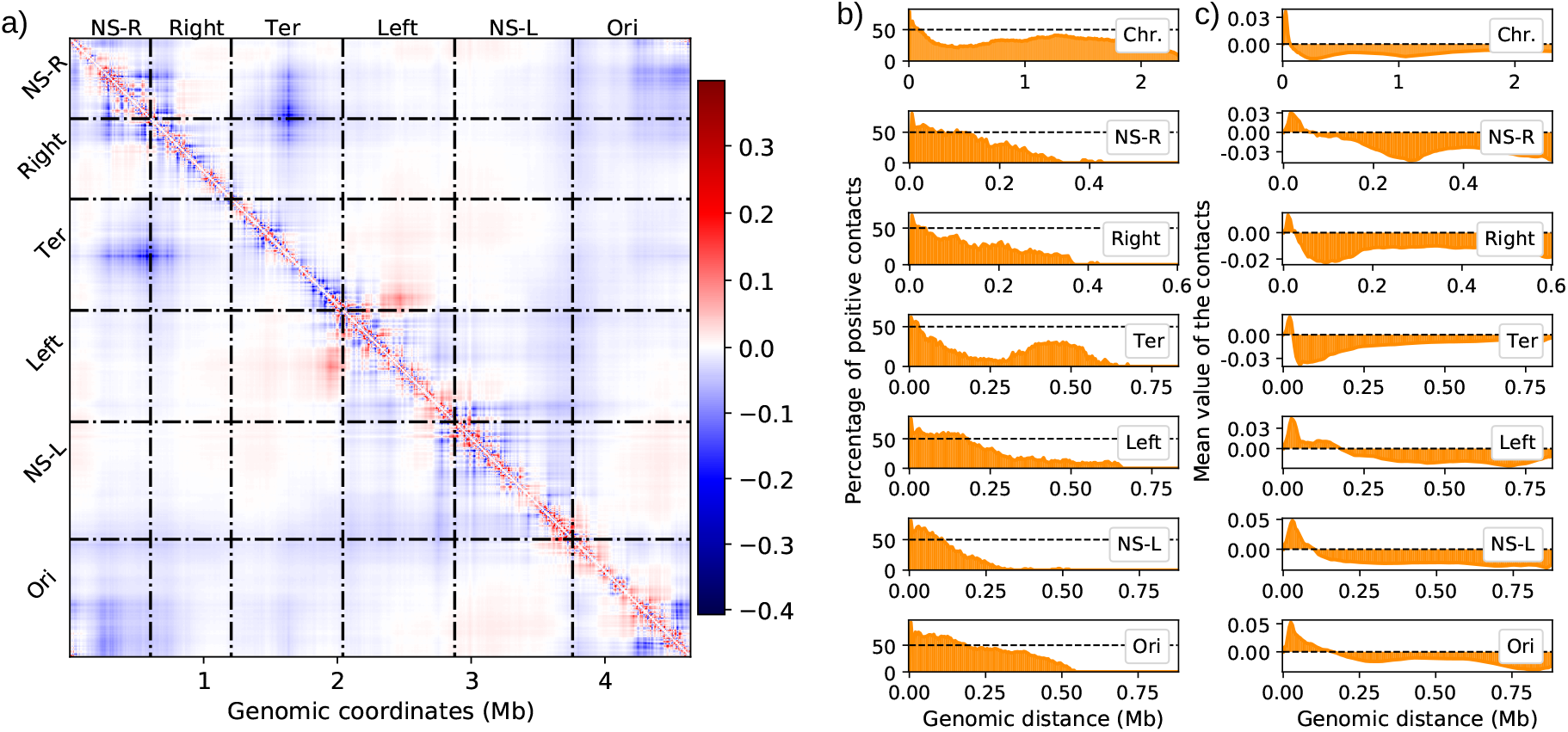
**a)** Difference heatmap for ΔhupAB mutant (ΔhupAB - WT), **b)** percentage of positive contacts from the difference heatmap with respect to genomic distance, **c)** average contact value from the difference heatmap with respect to genomic distance, for whole chromosome and macrodomains.

Furthermore, from the same difference heatmap, we have calculated the percentage of positive contacts, and average value of the contacts (for rational, see *SI methods*), shown in Figure 3b and c, respectively. If the percentage of positive contacts is greater than 50% and mean contact value is also positive i.e. > 0, it imply that the contact probability, between the loci separated by that particular genomic distance, on average has increased in case of mutant, otherwise decreased with respect to WT. Similar to Hi-C experiment, we also found that, in ΔhupAB mutant, the contact probability was increased on an average at shorter length scales for whole chromosome, as shown by the higher positive contacts and average contact value > 0 in Figure 3b and c, respectively. For individual macrodomains and non-structured regions also, we observed similar results (Figure 3b and c). In case of NS-R, Right and Ter the positive contacts show increase (i.e. > 50%) only for a very short genomic distance. Whereas for Left, NS-L and Ori, the average number of positive contacts (which are > 50%) sustained up to 100-200 Kb. In Ter, we observed an interesting trend in which, first the percentage of positive contacts decreased till the genomic distance of ~ 300 Kb and then increases till ~ 450 Kb, whereas for other MDs the percentage didn’t show significant increase. Therefore it seems that the effect of hupAB mutation is different in case of Ter than other macrodomains, indicating HU’s activity in Ter packing in the range of ~ 25 − 300 Kb, similar to what was found in experiment[23]. Altogether, these observations suggest the similarity between our simulation and experimental results, however, we were unable to find any distinct demarcation for contacts range through the contact difference heatmap, which was observed in the experiment[23].

These results from our simulations are consistent with experiments which hypothesized that HU is involved in long range interactions (> 280 Kb) in whole chromosome[23]. As observed in our simulations with the ΔhupAB mutant cell, when compared with WT *E. coli*, the long range interactions have diminished but local or short range interactions were enhanced or not affected significantly.

### FIS protein

FIS is one of the most abundant NAPs in rapidly growing *E. coli* cells [34]. Initially, FIS was considered mainly as a regulatory protein, involved in site-specific DNA recombination[35, 9]. Later on, studies showed that, based on FIS concentration, it can bind to DNA non-specifically also, and play a role in nucleoid organization[12, 13]. When we look at the ΔFIS mutant results, experimental and simulated matrices (Figure S4a, and S4b respectively) are in mutually good agreement (R = 0.92). At local scale (i.e. near the main diagonal) the respective WT (Figure 4a upper triangle) and ΔFIS mutant (Figure 4a lower triangle) are quite similar whereas long range contacts have diminished in the ΔFIS mutant. Similar results were obtained from the R_*g*_ map (Figure S2 c and d) in which the R_*g*_ for each segment has increased but the pattern is same as WT with high correlation coefficient of 0.84. However, unlike ΔhupAB mutant the correlation decreases faster for larger genomic segments (Table S5).

**Figure 4:**
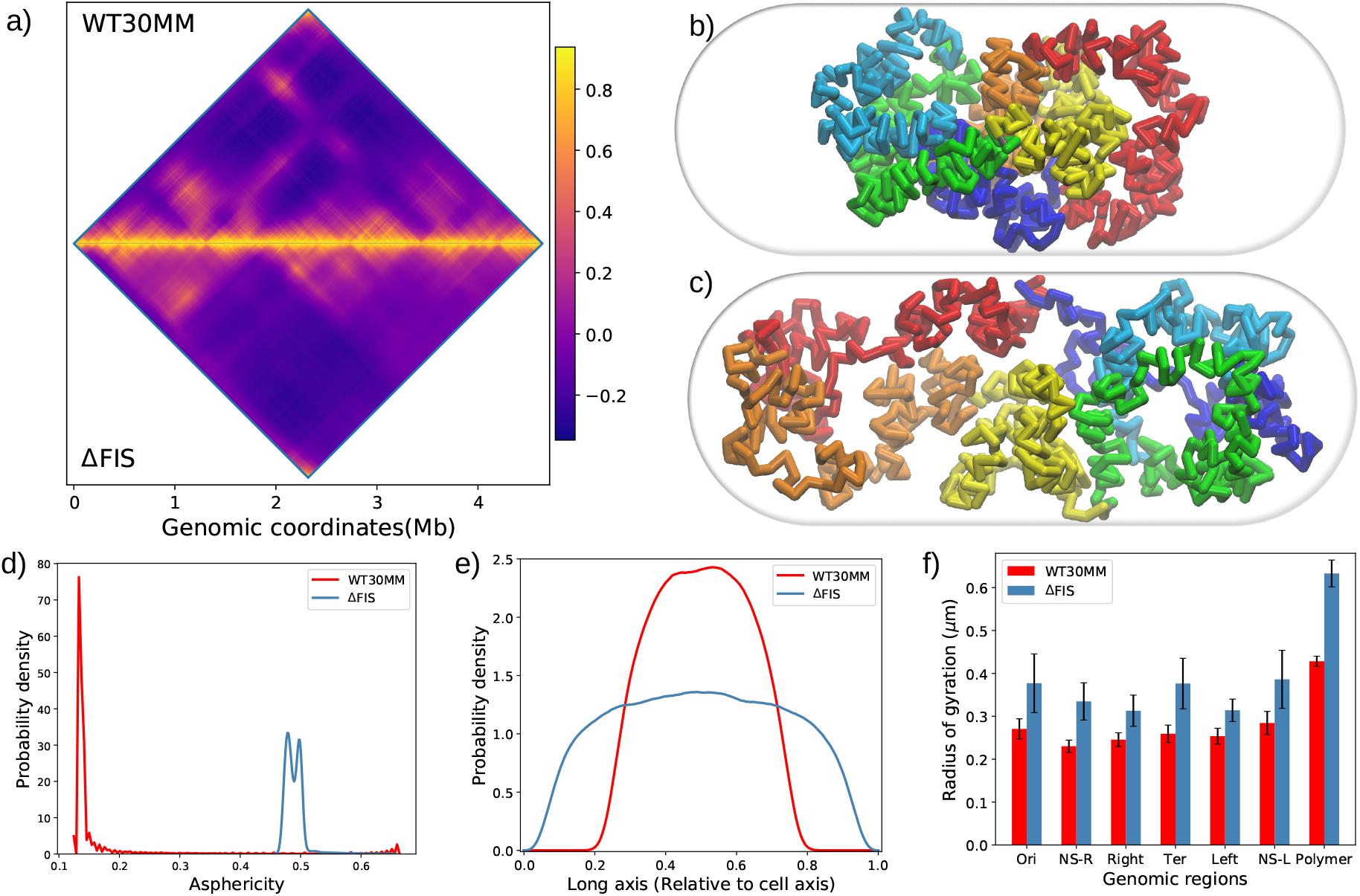
**a)** Correlation matrix of simulated WT (upper triangle) and ΔFIS (lower triangle) contact probability matrix, **b)** snapshot from equilibrated structures for WT *E. coli* (grown in MM at 30°C) chromosome simulation, **c)** snapshot from equilibrated structures for ΔFIS mutant chromosome simulation, **d)** distribution of asphericity of WT and ΔFIS mutant cells, **e)** linear DNA probability density for WT and ΔFIS mutant cells relative to cell length along the long axis (Z- axis), **f)** radius of gyration (R_*g*_) comparison between WT cells and ΔFIS mutant at same growth conditions.

In Hi-C experiment, the ΔFIS mutant showed contact enrichment in the 5-100 Kb range but our difference heatmap analysis between ΔFIS mutant and respective WT (Figure S5a) suggests no significant increase in contact probability in that range. As shown in Figure S5b, the percentage of positive contacts from difference heatmap is also very low (< 50%) along with negative average contact values (S5c) implying decreased contact probability in ΔFIS mutant. In case of Ter macrodomain also, our model does not predict any significant increase in contact probability below 100 Kb and there is a general decrease in contact probability throughout the chromosome and in individual macrodomains.

When we look at the chromosome structure, Figure 4c shows one of the representative conformations from simulations for the ΔFIS mutant cells, which is visibly more expanded than its corresponding WT chromosome at similar conditions (in MM at 30°*C*, Figure 4b). The relative change in asphericity and axial chromosome density, in comparison to WT as shown in Figure 4d and e respectively, also predict the relative expansion of the chromosome in ΔFIS mutants. R_*g*_ values, given in Figure 4f, suggest that each of the individual macrodomains are affected to the similar extent due to lack of FIS but the overall effect on whole chromosome is more severe than individual macrodomains. This is more prominently visible through their intra-domain rms end-to-end distance plot given in Figure S6.

Our results suggest qualitative similarity in the effect of ΔhupAB and ΔFIS mutations on bacterial chromosome structure. The distribution of axial density for each macrodomain is similar for both models as shown in Figure S7a and b, except Ter. Intra-domain distance profile, as shown in Figure S8, also indicate similarity between overall folding of macrodomains in both the mutants. However, the changes in whole chromosome conformation are more dramatic in case of ΔFIS mutant than ΔhupAB, with respect to their WT counterparts. The asphericity and axial density of model chromosomes for both the mutants are similar as given in table S2. When we compare these effect with their WT counterparts (*E. coli* at 37°*C* in LB and 30°*C* in MM for hupAB and ΔFIS mutant, respectively), the asphericity increases from 0.134 to 0.499 in case of ΔFIS whereas for ΔhupAB mutant it only increases from 0.374 to 0.503. Same is true for linear density as well. Since HU helps in sustaining negative supercoils in the chromosome along with gyrase (gyrB)[36] and similar role was found for FIS [37], it is possible that the effect of mutation of any of the two proteins will have similar effect, which we are observing in our current model as well.

As a part of dual role of FIS in gene regulation and chromosome organization, it is known to mediate formation of dense clusters of DNA at highly transcribed regions such as rrn operons[10]. Through insertion of fluorescent protein binding sites and epifluorescence microscopy, all rrn operons but rrnC were recently shown to co-localize and form a structure similar to eukaryotic nucleolus of the size of around 80 to 130nm[38]. On the other hand, deletion of nearby FIS binding sites resulted in disruption of cluster. To quantify the role of FIS on potential clustering of rrn operons, we calculated and compared average distance between rrn (between two rrn operons) and non-rrn pairs (having at least one non-rrn locus) for both WT and ΔFIS mutant as shown in Figure 5. In case of both WT and ΔFIS mutant, the spatial distance scaling with genomic distance is similar for rrn and non-rrn pairs, generally increasing with genomic distance. However the logarithmic scaling of the spatial distance indicates that for larger genomic distance the distance between two loci generally saturates.

**Figure 5:**
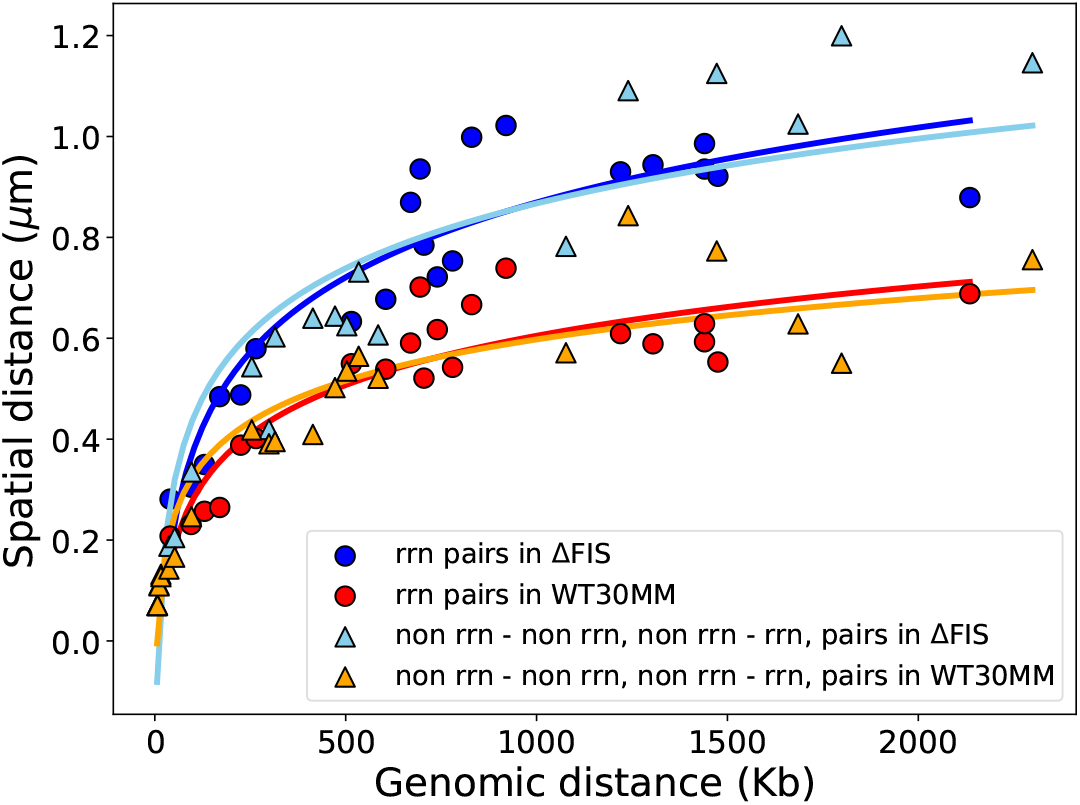
Average spatial distance with respect to genetic distance in rrn operon pairs (red circles for WT and blue circles for ΔFIS mutant) and non rrn - non rrn and non rrn - rrn pairs (orange triangles for WT and cyan triangles for ΔFIS mutant).

In the experiment, rrn pairs without rrnC showed co-localization, but based on the average distance between rrn operons (Table S3), we were unable to find any significant difference in distance between pairs with and without rrnC in both WT and ΔFIS mutant. In both the cases the distance increased with genomic distance. Also, the average distances in rrn operon pairs (Table S3) are larger than what was reported originally [38], ranging from 200 − 750nm in WT and ~ 300 − 1000nm in ΔFIS mutant. However, we did see a relative increase in the distance between the rrn operons (similarly in non-rrn pairs also) in the ΔFIS mutant but it could be due to the overall expansion of the chromosome, rather than any specific effect of ΔFIS mutation on the operons. The results are in accordance with the original Hi-C experiment[23] in which the contact frequency didn’t indicate any clustering in the rrn operons. In our simulations as well we weren’t able to find clustering of rrn operons. Altogether, our results also follow the similar trend as Hi-C experiment[23] and contradict the results of fluorescence study[38]. As suggested, to shed more light on the issue, more experiments are needed[39].

### MatP protein

MatP is a crucial protein for cell division and chromosome segregation. It binds specifically to the 23 sites (called as matS) in Ter macrodomain in the *E. coli* chromosome[17]. Currently, two working mechanisms have been proposed for Ter macrodomain insulation by MatP protein, based on structural studies[40, 41] and Hi-C experiment[23]. First is, by forming MatP tetramers and bridging distant matS sites[40], and second, by excluding MukBEF condensin from Ter macrodomain. Since MukBEF forms long range contacts, Ter interactions are limited to its own self[23, 41].

To determine the effect of MatP mutation on Ter macrodomain and whole chromosome, we performed similar simulations as in previous sections with Hi-C data from ΔMatP mutant. Processed Hi-C data details are given in table S1. Experimental and simulated contact probability matrices (Figure S9a and S9b respectively) are in reasonable mutual agreement with a correlation coefficient of 0.93. We see that in the ΔMatP mutant the contact probability is similar to the WT at smaller genomic distances, as shown in Figure 6a. Also, the local structure, explored by R_*g*_ map, shows similarity between WT and ΔMatP mutant, shown in Figure S2 c and e.

**Figure 6:**
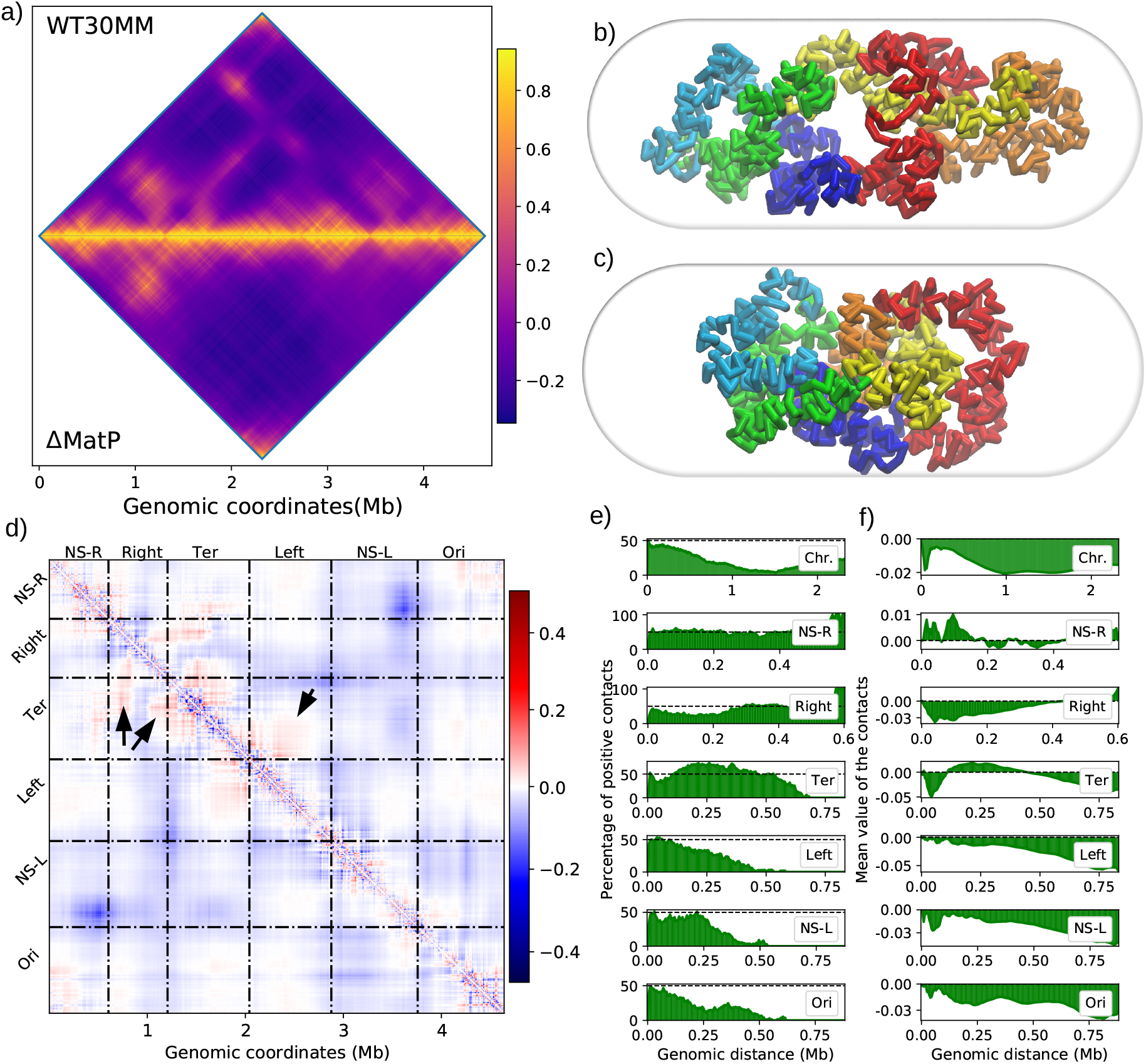
**a)** Correlation matrix of simulated WT (upper triangle) and ΔMatP (lower triangle) contact probability matrix, **b)** snapshot from equilibrated structures for ΔMatP mutant chromosome simulation, **c)** snapshot from equilibrated structures for WT *E. coli* (grown in MM at 30°C) chromosome simulation, **d)** difference heatmap for ΔMatP mutant (ΔMatP - WT30MM), **e)** percentage of positive contact, and **f)** average contact value, calculated from the difference heatmap for ΔMatP mutant.

Structurally, (Figure 6b and c), the chromosome has become more loosely packed in ΔMatP mutant than in WT cells. The spread of the overall chromosome can be seen from the axial density plot and asphericity comparison (Figure S10a and b). Similar results are obtained from radius of gyration of macrodomains and whole chromosome, shown in Figure S10c. Interestingly, the R_*g*_ has increased for only Left, NS-L, and Ori macrodomains whereas for NS-R, Right, and Ter it remained unchanged. Similarly in intra-domain distance plot (Figure S11), the spatial distance increased for the Left, NS-L and Ori for genomic distances larger than 100-200Kb, in ΔMatP mutant. In case of other three regions (NS-R, Right, and Ter) the results are varying. For NS-R the intra-domain distance is almost unchanged throughout whole genomic distance range. In Right macrodomain, the spatial distance is greater in ΔMatP mutant at small to intermediate genomic distances and then decreases. Whereas in Ter, the spatial distance increases at small genomic distance followed by decreasing in intermediate range and finally increasing for large genomic distances (> 500Kb).

Difference heatmap between ΔMatP mutant and WT (Figure 6d) indicates decreased long range interactions and enhanced short ranged ones, similar to previous mutants. But, percentage of positive contacts and average contact values, as shown in the first row of Figure 6e and f, shows overall decreased interaction at whole chromosome scale. The positive contacts are less than 50~ and the average contact value is also negative at all genomic distances. But interactions inside and in between macrodomains show different results. Since MatP is involved in Ter packing and isolation, its removal affected Ter and its flanking macrodomain differently than others. The contact probability of Ter within itself and with Right and Left macrodomains has enhanced in the ΔMatP mutant which is evident by the red/positive contact between Ter-Right and Ter-Left region, shown by the black arrows in Figure 6d. However, a significant increase in percentage of positive contacts and average contact value is only observed in Ter and Right macrodomains (third and fourth row in Figure 6e and f). These changes in Ter and its flanking macrodomains, especially Right, are in agreement with the previous experiment[17]. Inside Ter we also observed a decrease in short range contacts in ΔMatP mutant but only below 100 Kb after which the contacts were increased till ~ 450 Kb. These results for Ter also corroborate the results obtained from intra-domain distance plot (Figure S11). For NS-R, as shown in Figure 6e and f (second row), the percentage of positive contacts almost remains near 50~ throughout the genome except at very large genomic distance and the average contact value fluctuates around 0. Therefore the average effect of ΔMatP mutation on NS-R region is negligible which also explains the result obtained in intra-domain distance plot. Therefore in ΔMatP mutant, the individual macrodomains affected are Ter and Right, but interactions between Ter and Left are also enhanced without affecting Left macrodomain in itself. In other macrodomains the contact probability decreases with genomic distance indicating their expansion, as found by difference heatmap analysis (Figure 6d, e and f).

In the recombination assay experiment for MatP mutant, the interaction frequency between Right and Ter macrodomain was increased in contrast with wildtype, indicating Ter disorganization. Therefore for further investigation, we compared the distances between ter2 and ter4 loci, which are 100kb apart in Ter region (Figure 7a), we observed an average increase in the interlocus distance in ΔMatP mutant. These two loci show that the parts of Ter macrodomains are disorganized while keeping overall properties same. We also calculated the distance between ter2 and ter6 loci, which are ~ 350 kb apart (Figure 7b). We found that in wildtype the average distance is 0.466**μ**m whereas in ΔMatP mutant it decreased to 0.451**μ**m. But in case of wildtype the variance is very low implying greater rigidity in motion between these two loci. In ΔMatP mutants, though the average is lower but the variance is more than double from WT which indicates the flexibility and disorganization between these two loci. Therefore these results along with previous results suggest an apparent disorganization of Ter and flanking region without significant change in its overall structure.

**Figure 7:**
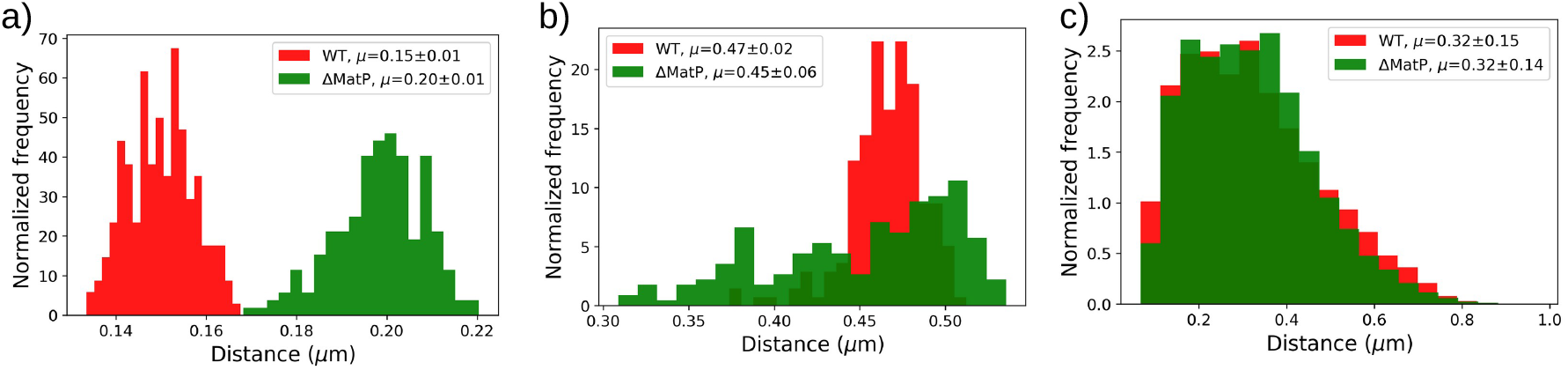
**a)** distance between ter2 and ter4 loci which are 100kb apart in Ter region in *E. coli* chromosome for WT and ΔMatP mutant cells. **b)** distance between ter2 and ter6 loci which are ~ 350kb apart in Ter region in *E. coli* chromosome for WT and ΔMatP mutant cells, **c)** distribution of distance between all 23 matS sites in the *E. coli* genome in WT and ΔMatP mutant cells.

Since there are two mechanisms proposed for the isolation of Ter macrodomain by MatP protein as explained earlier, we also investigated the possible mechanism from obtained structures. Similar to the Hi-C experimental results[23], our structures were also devoid of any significant changes in matS site’s contact probability/distance in wildtype and ΔMatP mutant cells. As shown in Figure 7c, the distribution of distance between all 23 matS sites showed only slight variation between WT and ΔMatP mutant. Therefore our results also support the argument by *Lioy et al.*[23] that the Ter organization doesn’t require MatP protein to bring the matS sites in close proximity. And altogether we find that in ΔMatP mutants, most of the chromosomal properties remain conserved with decrease in long range interactions throughout whole chromosome except Ter. Ter and its flanking regions showed increased long range interaction along with decrease in short ones.

### MukBEF protein

In *E. coli* there is only single SMC complex present, MukBEF, which is suggested to play a major role in Ori macrodomain positioning and chromosome segregation. It is composed of 3 subunits, MukB, MukE, and MukF, in which MukB has the DNA binding property and MukE and MukF stabilizes the complex[21]. It was first identified as a protein crucial for chromosome segregation in *E. coli* cells and required for correct positioning of Ori[18, 19, 20]. Also, it is known to facilitate DNA loops formation and helps in its compaction[21]. Therefore MukBEF has shown to play an important role in DNA organization and compaction and thus it is an indispensable part of the *E. coli* nucleoid. After integrating the experimental Hi-C matrix for MukB mutant (ΔMukB *E. coli* in MM at 22°C, Figure S12a) into our model, we were able to recover the simulated contact probability matrix (Figure S12b) with a Pearson correlation coefficient of 0.93. Comparison with simulated WT matrix (Figure 8a, upper triangle), the ΔMukB mutant suggests decreased long range interactions in the ΔMukB mutant (Figure 8a, lower triangle).

**Figure 8:**
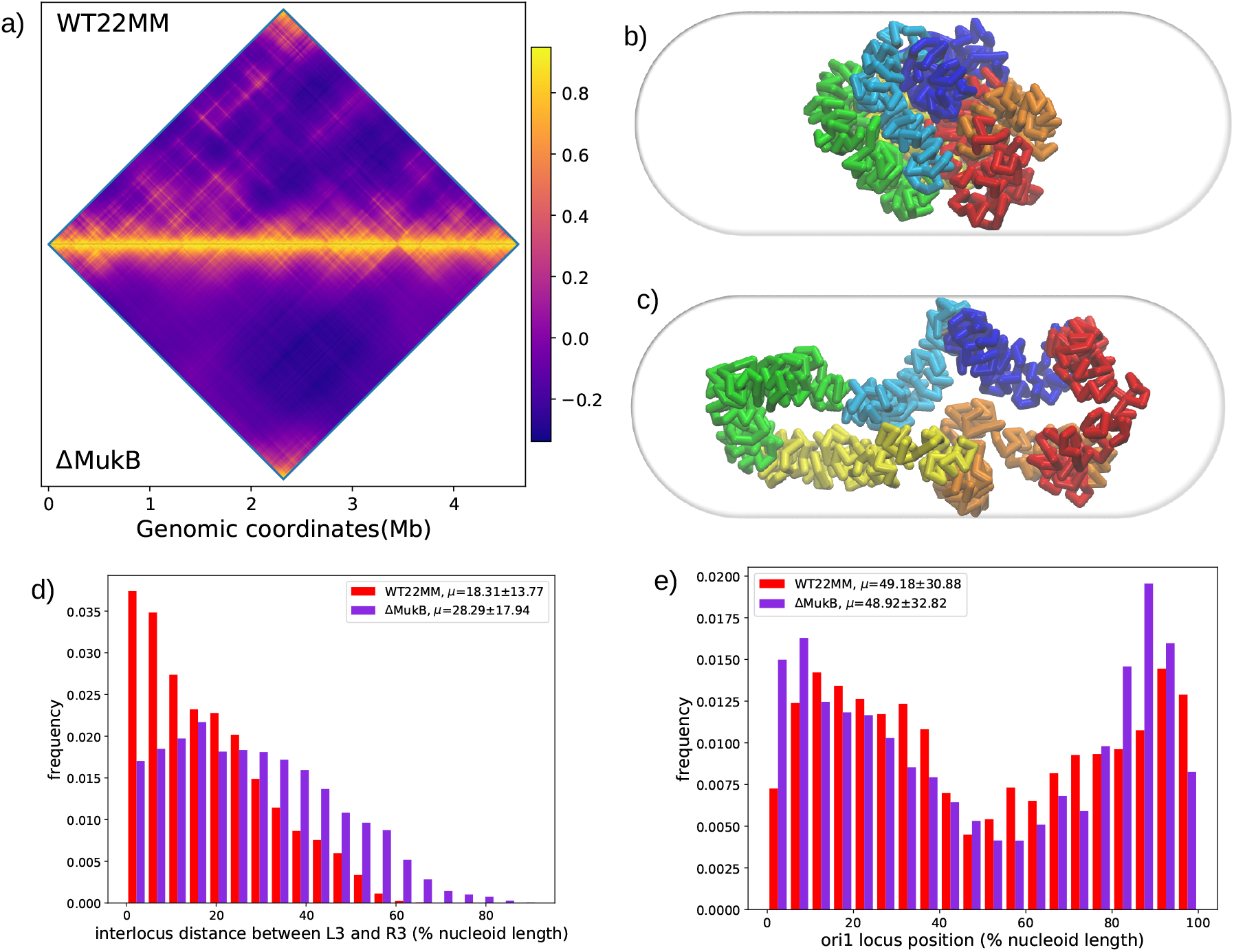
**a)** Correlation matrix of simulated WT (upper triangle) and ΔMukB (lower triangle) contact probability matrix, **b)** snapshot from equilibrated structures for WT *E. coli* (grown in MM at 22°C) chromosome simulation, **c)** snapshot from equilibrated structures for ΔMukB mutant chromosome simulation, **d)** L3-R3 interlocus distance distribution in WT (red) and ΔMukB mutant (purple), **e)** Ori1 position distribution in WT (red) and ΔMukB mutant (purple).

Structurally, the chromosome appears to be expanded along the axial dimension in the ΔMukB mutant with respect to the WT (Figure 8c and b, respectively). Asphericity plot and axial density profile (Figure S13b and c) also suggests the same as both have increased in ΔMukB mutant. The radius of gyration for whole chromosome, as show in Figure S13c, also indicates the expansion of chromosome along with all the macrodomains and NS regions except Ter. In Ter the effect of ΔMukB mutation was minimal. R_*g*_ map from ΔMukB mutant revealed similar results to previous mutants i.e. similarity with its WT counterpart(Pearson’s correlation coefficient = 0.84, at 100 Kb window size, Figure S2f and g).

Difference heatmap (Figure S14a) also showed similar results. Whole heatmap is largely occupied by the negative/blue contacts indicating decreased long range contacts but near the diagonal positive/red contacts are also present suggesting enhanced short or intermediate ranged contacts. Furthermore, the percentage of positive contacts and average contact values from the difference heatmap (Figure S14b) shows an increase in short to intermediate range contacts on a chromosome as well as individual macrodomain scale. Therefore, it seems that the effect of ΔMukB mutation was similar in all macrodomains.

Although, the role of MukBEF is more pronounced in replicating cells but in non-replicating cells, absence of MukBEF showed mispositioning of Left, Right and Ori macrodomains[42]. Therefore, we also determined the positions of L3, R3, and ori1 loci (representative loci for Left, Right, and Ori macrodomain) and the interlocus distance between L3-R3, as previously explored[42]. However, our results lack the similarity with those experimental results. There was a ~ 10% increase in the distance between L3-R3 loci, as shown in Figure 8d, in the ΔMukB mutant, whereas in the absence of MukE or MukF the distance was decreased in the experiment. Also, the percentage of L3-R3 loci being in same nucleoid half was decreased from 46.78% in WT to 27.46% in ΔMukB mutant, contrary to the experiment, where after removing MukBEF the percentage was increased[42]. Position of ori1 loci, which is a representative of the Ori macrodomain (Figure 8e), showed negligible change in ΔMukB mutant. However, experimentally the ori1 loci moved towards the edge of the nucleoid in the MukE or MukF absence, in our case the ori1 is showing preference for the nucleoid edge in WT as well as ΔMukB mutant, as shown in Figure 8e. To gain a better picture of macrodomain arrangement in WT and mutant we also calculated the axial density distribution of all macrodomains and NS regions as shown in Figure S15a and b, for WT and ΔMukB mutant, respectively. From the axial density of macrodomains it seems that in case of ΔMukB mutant, the separation between macrodomains has increased and they are well distributed in the nucleoid. Apart from positions, the fluctuations in the position, as calculated by standard deviations, has also increased mainly in case of Ori and Ter. Therefore altogether, due to expansion of whole chromosome the macrodomains show better separation along axial axis and increased fluctuations in the mean position of Ter and Ori macrodomains.

Since MukBEF is known to interact with MatP and affect Ter organization[41, 23], we have also calculated the interlocus distance between ter2 - ter4, and ter2 - ter6 for ΔMukB mutant (Figure S16a and b, respectively). In ΔMukB mutant the distance between ter2 and ter4 appears to be decreased (Figure S16a), whereas it increased in case of ter2 and ter6. These results are consistent with the observations of difference heatmap analysis, since the contact probability was enhanced in ΔMukB mutant for short ranged interactions, the distance between ter2 and ter4 loci (100 Kb apart) was expected to decrease. Probability for long range interactions were decreased therefore distance between ter2 and ter6 was increased. Similar to the ΔMatP mutant results, distribution of distance between matS sites pairs are similar in WT and ΔMukB mutant also (Figure S16c). Therefore in ΔMukB mutant the Ter macrodomain appears to be affected as the distance between the loci was increased but matS sites distribution lack significant changes.

## Discussion

The current study provides a quantitative basis of the stabilization effects of various nucleoid associated proteins (NAPs) on *E. coli* chromosome organization. We attempted to utilize a recently devised simulation method [22] and combined the Hi-C data of NAPs mutants (individually) with a beads-on-a-spring model and compared the results with their respective wildtype (WT) simulations. We have studied the effect of 3 NAPs HU, FIS, MatP, and a condensin protein MukBEF and the result of each of the mutations, on overall chromosome organization, are summarized in Figure 9. As with the WT, we were able to reproduce the experimental Hi-C interaction matrix for the mutants with good correlation as shown in table S1. HU and FIS are know to bind throughout whole chromosome and organize it at large genomic scale[9, 13, 23]. MatP binds in Ter macrodomain, specifically at 23 MatS sites and maintain Ter organization. MukBEF is a condensin like protein which plays major role in daughter chromosome segregation and maintain Left-Ori-Right configuration of the chromosome in *E. coli* [42].

**Figure 9:**
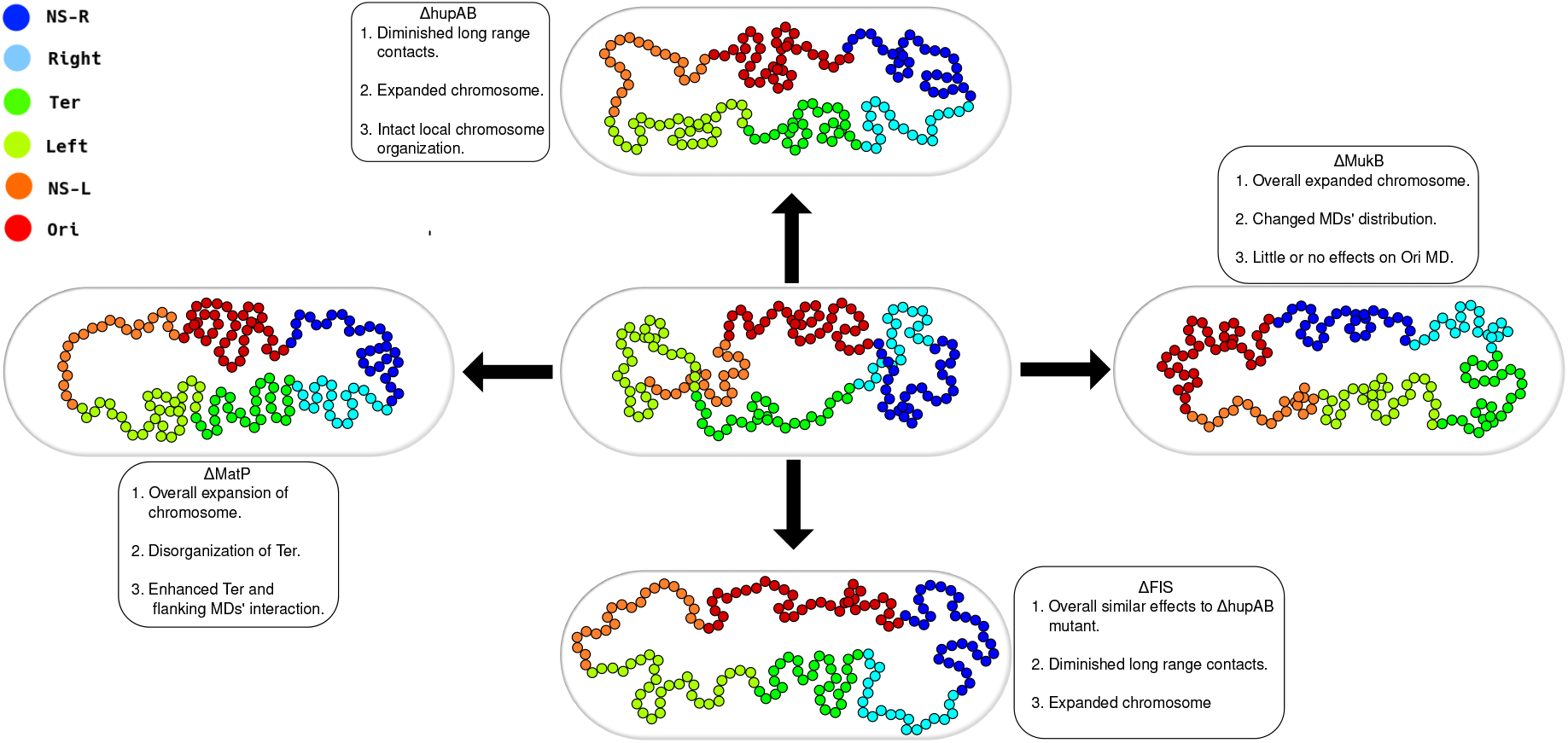
**a)** Schematic for the WT, **b)** ΔhupAB mutant, **c)** ΔFIS mutant, **d)** ΔMatP mutant, **e)** ΔMukB mutant *E. coli* chromosome.

In case of ΔhupAB mutant we observed that the long-range contacts were diminished which is apparent from the simulated contact probability matrix itself. However, the difference heatmap analysis revealed an average increase in short range contacts, the reason could be decrease in the stiffness of the DNA. As previously shown, excess of HU can lead to formation of filamentous DNA with increased stiffness[43], which can hinder the local chromosome interactions. Therefore, deletion of hupAB from the cells may have led to increased short ranged contacts. From the structure of chromosomes in 200 trajectories, the chromosome seems to be getting decondensed and occupying whole cell volume in ΔhupAB mutant. The expansion of chromosome is also evident from the asphericity, linear density and radius of gyration (R_*g*_) of the chromosome. The asphericity, which indicates the extent of sphericity in the structure[29, 44], predicts more cylindrical structure for ΔhupAB mutant chromosome. Linear density and R_*g*_, both indicate an overall expanded structure. Individual macrodomains and non-structured regions also showed an increase in their R_*g*_. However, the change in case of Ter was minimum as observed in Hi-C experiment itself.

ΔFIS mutants showed similar results to that of ΔHU mutants. There was an overall increase in long range contacts with minute increase in short range ones as determined by difference heatmap analysis. Whereas R_*g*_ indicates an overall chromosome expansion, asphericity and linear density indicate expansion in axial dimension, along the cell length. However, when compared with their respective wildtype counterparts, FIS deletion showed much more drastic changes than hupAB deletion. The macrodomains are also more expanded in ΔFIS mutant than ΔhupAB evident by intra-domain distance plot. Another interesting feature of FIS protein is its role in formation of nucleolus-like structure by rrn operons in *E. coli* [38]. But when we computed the average distance between all rrn operon pairs from the wildtype *E. coli* itself, we did not observe any co-localization and the average distance also varied extensively. Although the average distance was increased in case of FIS mutant, it was mainly due to overall chromosome expansion because the distance between non-rrn operon pairs has also increased to a similar extent. Also, the signature of co-localization was absent in original Hi-C interaction matrices, which further supports our results.

MatP is important for chromosome positioning and segregation during cell division. When we simulated chromosome for MatP mutant cell, we observed an overall chromosome expansion but individual macrodomain properties remained similar to wildtype. The interactions between Ter and its flanking macrodomains were enhanced in ΔMatP mutant along with Ter disorganization as the distance between two loci, in Ter region (ter2 and ter4, 100Kb apart), was increased in the MatP mutant. Also, loci which were ~ 350 Kb apart (ter2 and ter6), showed increased variance in distance between them, indicating their disorganization in matP mutant cells Finally, the matS sites where MatP protein binds showed very less variation than wildtype, similar to what was observed in Hi-C experiment.

In ΔMukB mutants also the chromosome was expanded along the axial axis. The long range contacts were diminished and short- and intermediate- ranged contacts were enhanced. But in case of ΔMukB mutant the short ranged contacts were enhanced to a greater extent than any other mutant, shown by the difference heatmap analysis. We also observed that the position of ori1 locus was similar in both WT and ΔMukB mutant. Similarly, L3-R3 interlocus distance also, didn’t show significant change in the ΔMukB mutant. However, the linear density of the macrodomains, with respect to nucleoid length, have changed drastically in ΔMukB mutant. The linear density for both NS-R and Right have shifted towards right which is consistent with the experimental observation[23]. The Ter macrodomain got disorganized and its linear density is much more spread-out in case of ΔMukB mutant. For Ori and Left the changes are minor but NS-L density has shifted towards the left. Therefore, from the linear density we can clearly see how the distribution of macrodomains have changed in ΔMukB mutant. Also, it seems that relying on the position of a single locus in a macrodomain can be misleading for determination of that macrodomain position.

An interesting feature that we noticed in most of the mutant and WT samples was strong correlation among R_*g*_ maps, shown in Figure S17. For smaller regions i.e. at 50-100 Kb (10, 20 beads) the correlation between R_*g*_ map is high for all the mutants and wildtype with a net increase from 50 Kb to 100 Kb except for HU37-MatP30 and WT30-HU37 pair. However, for very small genomic distance (50 Kb, 10 beads) the correlation is slightly lower which could be due to stochastic nature of the interactions. Gradually, it decreases for larger segments starting from window size of 50 beads. However, the correlation between HU37 and FIS30 is almost constant for all window sizes, supporting our claim that the effect of FIS and hupAB deletions are similar. The overall comparison suggests that in all mutants and wildtypes the local chromosome packing remains similar which is expected because chromosome organization is largely regulated by transcription. For a cell to survive, expression of essential genes is required, which is going to be same in all cells. Therefore at scale of 50-100 Kb the overall structure of chromosome is same but at larger scale activity of proteins such as hupAB, FIS, MatP, and MukBEF and growth conditions play a major role.

Although, NAPs’ binding sites are a few base pairs long, these sites can be distributed throughout the chromosome, like for HU, FIS and MukBEF[45, 23, 1], or can be specific to some macrodomain, which is the case for MatP[23, 17]. But the current model is limited by the resolution of Hi-C interaction matrices and hence consider a coarse graining of 5000 bp to 1 bead, where multiple binding sites can be present in one coarse grained bead. This leads to our model being unable to answer questions which are specific to the role of NAPs in gene regulation or the change in the local structure of the chromosome due to the mutations. However, the current work has been largely successful in determining the effect of mutants on the overall structure of the chromosome.

