## Supporting information for "Nucleoid associated proteins and their effect on *E. coli* chromosome"

### Supplementary Information

#### Difference heatmap statistics

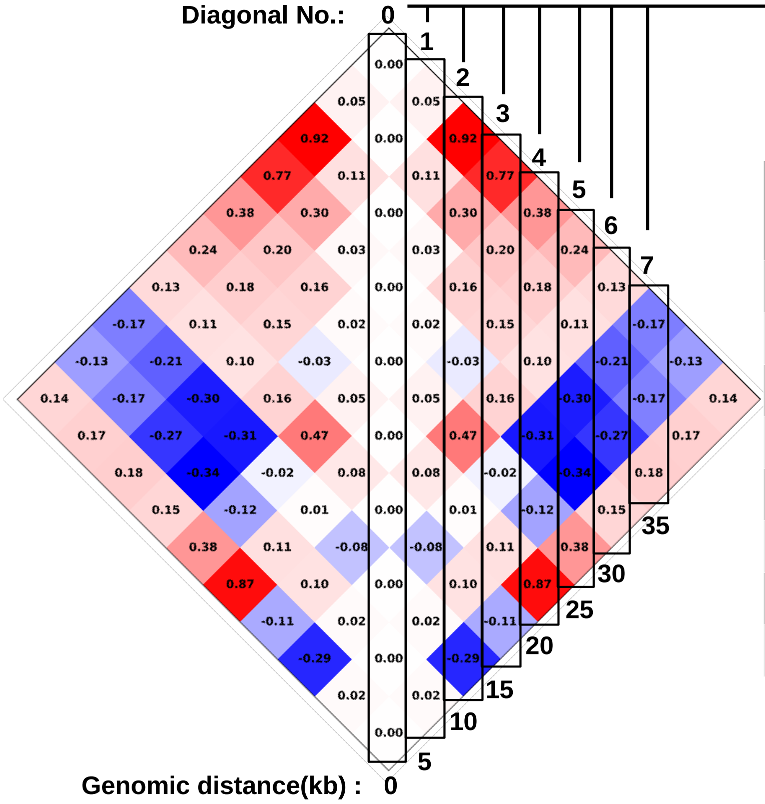


| Diagonal No. | Genomic Distance | % of positive contacts | Mean contact value |
|---|---|---|---|
| 0 | 0 | 0.00 | 0.00 |
| 1 | 5 | 88.89 | 0.03 |
| 2 | 10 | 75.00 | 0.21 |
| 3 | 15 | 71.43 | 0.18 |
| 4 | 20 | 66.67 | 0.18 |
| 5 | 25 | 60.00 | 0.02 |
| 6 | 30 | 50.00 | -0.05 |
| 7 | 35 | 33.33 | -0.06 |

**Figure 1:** Method for calculation of precentage of positive contacts and average diagonal value from the difference heatmap.

From the difference heatmap, which is calculated by subtracting WT matrix from the mutant, we have determined the percentage of positive contacts and mean contact value. These two properties provide us with an estimate of average changes in the contacts in mutants simulation with respect to wildtype simulation. As shown in Figure 1, we have selected the subsequent diagonals of the marix, numbered from 0 to n, where n is the length of the matrix. These diagonals, represent the contact values at a particular genomic distance, for example 5, 10, 15,... kb as shown with matrix in Figure 1. From these diagonals we have calculated the percentage of contacts which have positive values, for example, in $2^{nd}$ diagonal, out of 8 values 6 are positive and 2 are negative therefore the percentage of positive contacts for the diagonal will be 75% as shown in the table at the right. And similarly, from the same diagonal we will calculate the mean value for that particular diagonal which is 0.21 for the $2^{nd}$ diagonal. Now in this way we will calculate the percentages and average values for all the diagonals which can be plotted as a histogram. This whole process can pe performed for individual macrodomains also by selecting a square matrix for each macrodomain. We have calculated both the values because it might happen that there could be high number of positive contacts but a near zero average which indicate no significant change in contacts. Therefore to get a whole picture of the changes in contacts we have calculated the two properties.

### Supplementary Tables

**Table S1:** Hi-C processing details

| Mutant | Temperature (°C) | Media | No. of non-zero contacts | No. of contacts for simulation | Correlation coefficient(R) |
|---|---|---|---|---|---|
| WT | 37 | LB | 338939 (39.36%) | 17162 (5.06%) | 0.89 |
| WT | 30 | MM | 316148 (36.71%) | 28515 (9.02%) | 0.93 |
| WT | 22 | MM | 365406 (42.43%) | 46618 (12.75%) | 0.89 |
| ΔhupAB | 37 | LB | 247663 (28.76%) | 19972 (8.06%) | 0.95 |
| ΔFIS | 30 | MM | 219394 (25.48%) | 11619 (5.30%) | 0.92 |
| ΔMatP | 30 | MM | 350068 (40.65%) | 27118 (7.75%) | 0.93 |
| ΔMukB | 22 | MM | 267700 (31.08%) | 33059 (12.35%) | 0.93 |

**Table S2:** FWHM of linear density and Asphericity for all the samples

| Mutant | Temperature (°C) | Media | FWHM | Asphericity |
|---|---|---|---|---|
| WT | 37 | LB | 0.639 | 0.374±0.003 |
| WT | 30 | MM | 0.434 | 0.134±0.002 |
| WT | 22 | MM | 0.338 | 0.019±0.000 |
| ΔhupAB | 37 | LB | 0.795 | 0.503±0.003 |
| ΔFIS | 30 | MM | 0.800 | 0.499±0.001 |
| ΔMatP | 30 | MM | 0.631 | 0.386±0.003 |
| ΔMukB | 22 | MM | 0.553 | 0.314±0.003 |

**Table S3:** Mean and std of rrn operon pairs' distance (in $\mu m$) rounded off to nearest bead on the chromosome.

| rrn operon pairs | Genomic dist.(kb) | WT (in $\mu m$) | $\Delta$FIS (in $\mu m$) |
|---|---|---|---|
| rrnA-rrnB | 130 | 0.2570±0.016 | 0.3497±0.035 |
| rrnA-rrnC | 95 | 0.2309±0.008 | 0.3048±0.024 |
| rrnA-rrnD | 605 | 0.5387±0.127 | 0.6773±0.240 |
| rrnA-rrnE | 170 | 0.2646±0.027 | 0.4843±0.081 |
| rrnA-rrnG | 1305 | 0.5890±0.158 | 0.9439±0.384 |
| rrnA-rrnH | 830 | 0.6667±0.129 | 0.9985±0.336 |
| rrnB-rrnC | 225 | 0.3881±0.039 | 0.4877±0.096 |
| rrnB-rrnD | 740 | 0.6172±0.146 | 0.7216±0.288 |
| rrnB-rrnE | 40 | 0.2078±0.011 | 0.2811±0.022 |
| rrnB-rrnG | 1440 | 0.6289±0.166 | 0.9354±0.409 |
| rrnB-rrnH | 695 | 0.7015±0.130 | 0.9355±0.302 |
| rrnC-rrnD | 515 | 0.5495±0.127 | 0.6332±0.180 |
| rrnC-rrnE | 265 | 0.4012±0.051 | 0.5793±0.145 |
| rrnC-rrnG | 1220 | 0.6092±0.157 | 0.9299±0.369 |
| rrnC-rrnH | 920 | 0.7389±0.138 | 1.0218±0.365 |
| rrnD-rrnE | 780 | 0.5424±0.131 | 0.7533±0.305 |
| rrnD-rrnG | 705 | 0.5210±0.099 | 0.7846±0.227 |
| rrnD-rrnH | 1440 | 0.5930±0.153 | 0.9860±0.400 |
| rrnE-rrnG | 1475 | 0.5529±0.149 | 0.9211±0.425 |
| rrnE-rrnH | 670 | 0.5906±0.099 | 0.8694±0.256 |
| rrnG-rrnH | 2135 | 0.6883±0.207 | 0.8789±0.378 |

**Table S4:** Average rms end-to-end distance ($\mu$m) for the WT30MM and $\Delta$MatP mutant with respect to each macrodomain and non-structured regions.

| Mutant | Temperature ((°C)) | Medium | NS-R | Right | Ter | Left | NS-L | Ori | MatS | Rt-Ter |
|---|---|---|---|---|---|---|---|---|---|---|
| WT | 30 | MM | 0.563±0.005 | 0.565±0.004 | 0.453±0.005 | 0.452±0.004 | 0.510±0.004 | 0.515±0.006 | 0.623±0.005 | 0.565±0.005 |
| $\Delta$MatP | 30 | MM | 0.560±0.008 | 0.460±0.004 | 0.574±0.005 | 0.725±0.007 | 0.729±0.007 | 0.639±0.007 | 0.660±0.006 | 0.533±0.004 |

**Table S5:** Values for the Pearson's correlation coefficients between $R_g$ maps for all the WT and mutants pairs at various genomic distances (50, 100, 250, 500, 1000 kb) or bead window sizes (10, 20, 50, 100, 200). The correlations are reported for pairs, where, WT37: wildtype *E. coli* at 37°C in LB, WT30: wildtype *E. coli* at 30°C in MM, HU37: ΔhupAB mutant *E. coli* at 37°C in LB, FIS30: ΔFIS mutant *E. coli* at 30°C in MM, ΔMatP30: MatP mutant *E. coli* at 30°C in MM, and ΔMukB22: MukB mutant *E. coli* at 22°C in MM.

| Pairs | 50 | 100 | 250 | 500 | 1000 |
|---|---|---|---|---|---|
| WT37 - WT30 | 0.7065 | 0.7857 | 0.6812 | 0.4531 | 0.1074 |
| WT37 - WT22 | 0.5734 | 0.6968 | 0.6248 | 0.6251 | 0.5154 |
| WT37 - HU37 | 0.7693 | 0.8028 | 0.814 | 0.827 | 0.3827 |
| WT37 - FIS30 | 0.7563 | 0.8457 | 0.7875 | 0.6003 | 0.0397 |
| WT37 - MatP30 | 0.751 | 0.8406 | 0.6886 | 0.4105 | -0.1724 |
| WT37 - MukB22 | 0.5252 | 0.7279 | 0.6736 | 0.3897 | 0.11 |
| WT30 - WT22 | 0.6725 | 0.8468 | 0.8576 | 0.6607 | 0.4564 |
| WT30 - HU37 | 0.5753 | 0.5441 | 0.5059 | 0.4687 | -0.0302 |
| WT30 - FIS30 | 0.7235 | 0.8355 | 0.8086 | 0.6027 | 0.275 |
| WT30 - MatP30 | 0.8183 | 0.9059 | 0.9075 | 0.6682 | 0.1702 |
| WT30 - MukB22 | 0.6241 | 0.83 | 0.8791 | 0.7588 | 0.1517 |
| WT22 - HU37 | 0.4596 | 0.4537 | 0.4459 | 0.4861 | -0.2138 |
| WT22 - FIS30 | 0.5064 | 0.7058 | 0.7075 | 0.4889 | -0.1371 |
| WT22 - MatP30 | 0.6562 | 0.7907 | 0.8368 | 0.5627 | 0.3821 |
| WT22 - MukB22 | 0.6687 | 0.8427 | 0.8559 | 0.5971 | 0.2726 |
| HU37 - FIS30 | 0.6851 | 0.7266 | 0.6937 | 0.7 | 0.6428 |
| HU37 - MatP30 | 0.6449 | 0.6538 | 0.4677 | 0.2394 | -0.5901 |
| HU37 - MukB22 | 0.4031 | 0.4673 | 0.474 | 0.3649 | -0.1898 |
| FIS30 - MatP30 | 0.7517 | 0.8509 | 0.7909 | 0.5454 | -0.1576 |
| FIS30 - MukB22 | 0.4763 | 0.7085 | 0.7025 | 0.5506 | -0.1607 |
| MatP30 - MukB22 | 0.6164 | 0.8069 | 0.8792 | 0.8293 | 0.4724 |

### Supplementary Figures

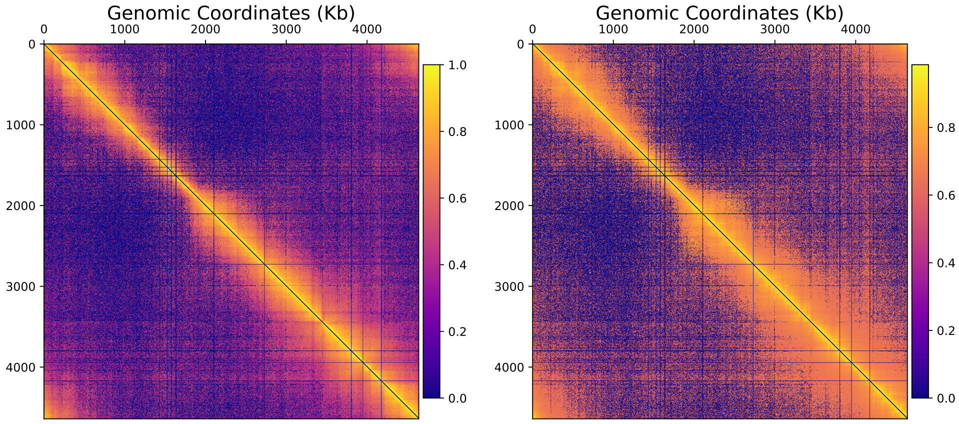


**(a)** Experimental matrix

**(b)** Simulated matrix

**Figure S1:** **a)** experimental contact probability matrix for ΔhupAB mutant *E. coli* MG1655 in LB at $37°C$, **b)** simulated average contact probability matrix for ΔhupAB mutant *E. coli* MG1655 in LB at $37°C$.

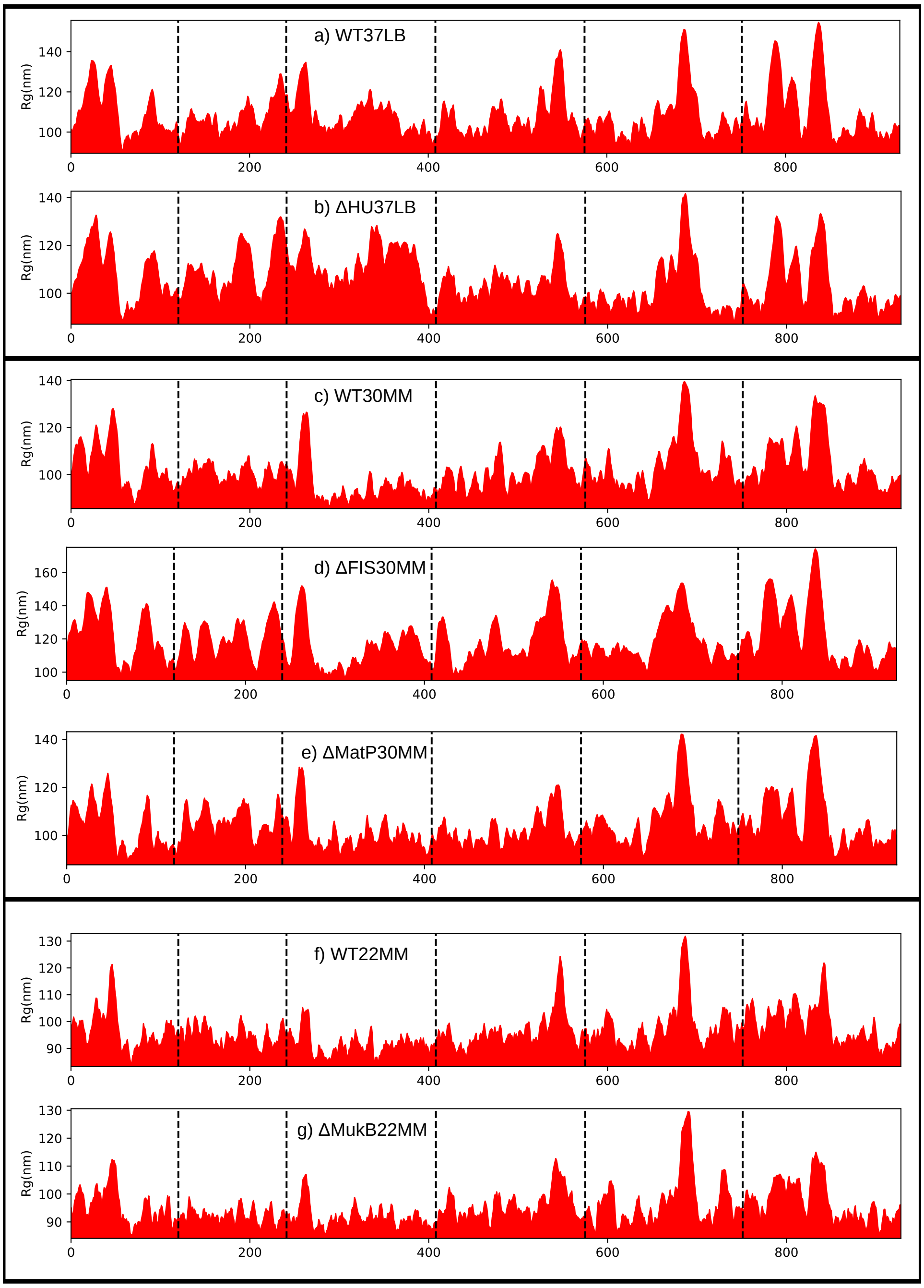
a) WT37LB
b) ΔHU37LB
c) WT30MM
d) ΔFIS30MM
e) ΔMatP30MM
f) WT22MM
g) ΔMukB22MM
Rg(nm)
0
200
400
600
800

**Figure S2**

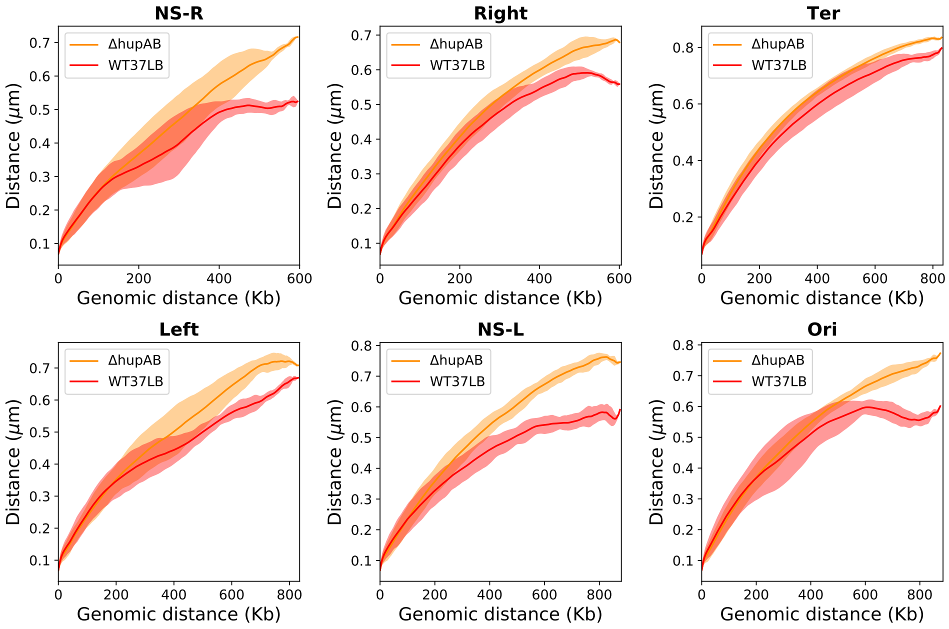


**Figure S3:** Rms end-to-end distance vs. genomic distance plot for intra-domain distances in ΔhupAB mutant (orange) and WT cells at 37°$C$ in LB media (red).

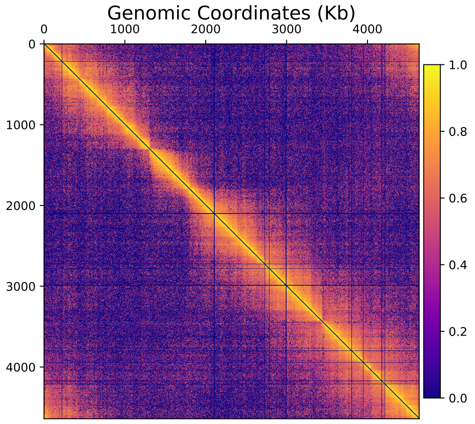


**(a)** Experimental matrix

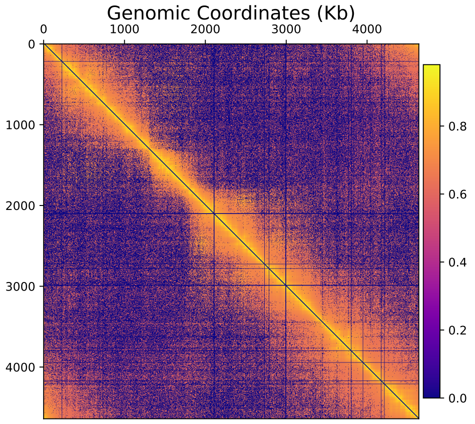


**(b)** Simulated matrix

**Figure S4:** **a)** experimental contact probability matrix for ΔFIS mutant *E. coli* MG1655 in MM at 30°$C$, **b)** simulated average contact probability matrix for ΔFIS mutant *E. coli* MG1655 in MM at 30°$C$.

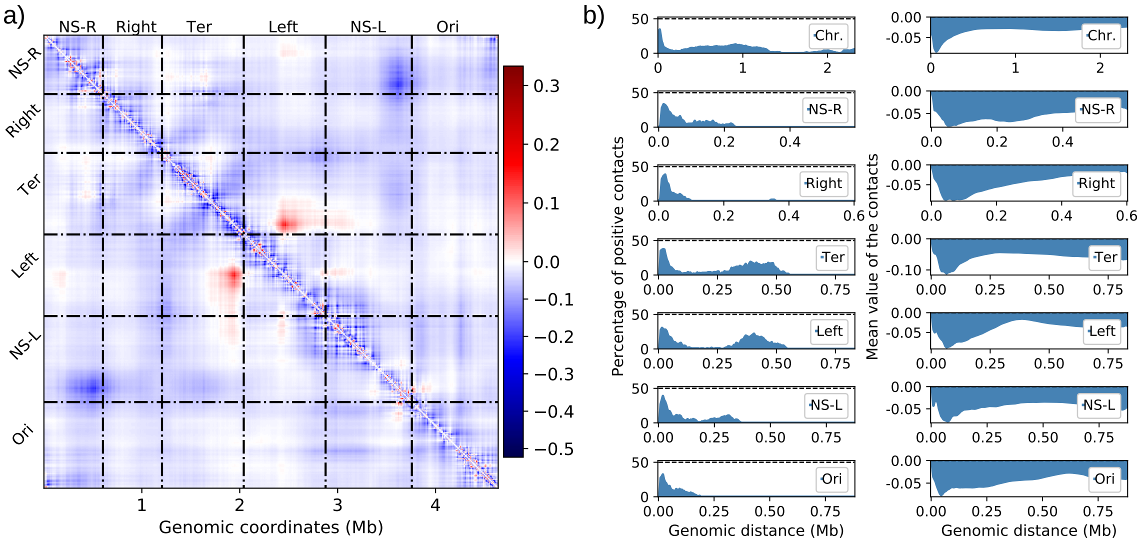


**Figure S5: a)** Difference heatmap for ΔFIS mutant (ΔFIS - WT30MM), **b) left** column: percentage of positive contacts from the difference heatmap (between ΔFIS - WT30MM) with respect to genomic distance, **right** column: average contact value from the difference heatmap (between ΔFIS - WT30MM) with respect to genomic distance.

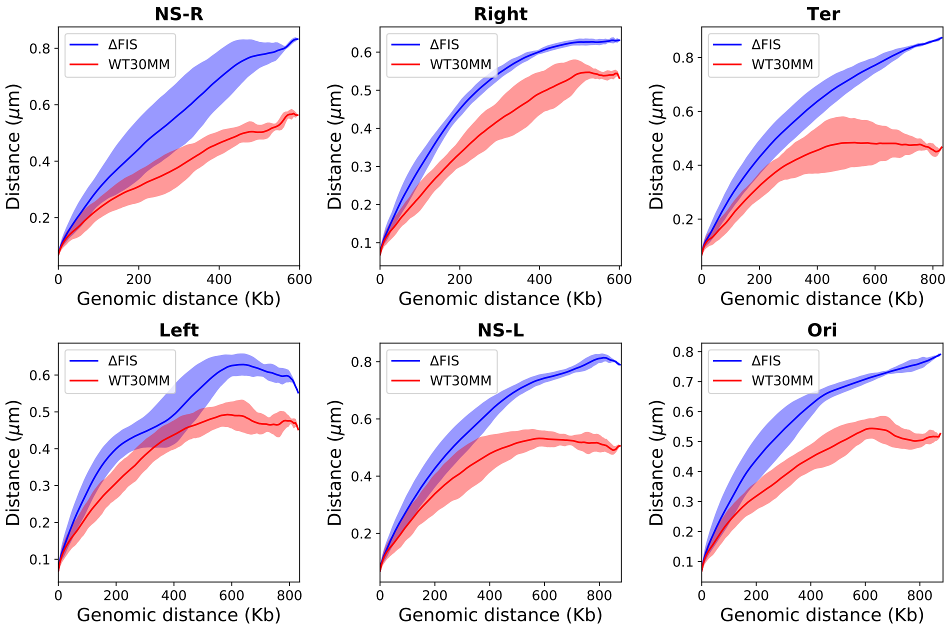


**Figure S6:** Rms end-to-end distance vs. genomic distance plot for intra-domain distances in FIS mutant (blue) and WT cells at 30°$C$ in MM media (red).

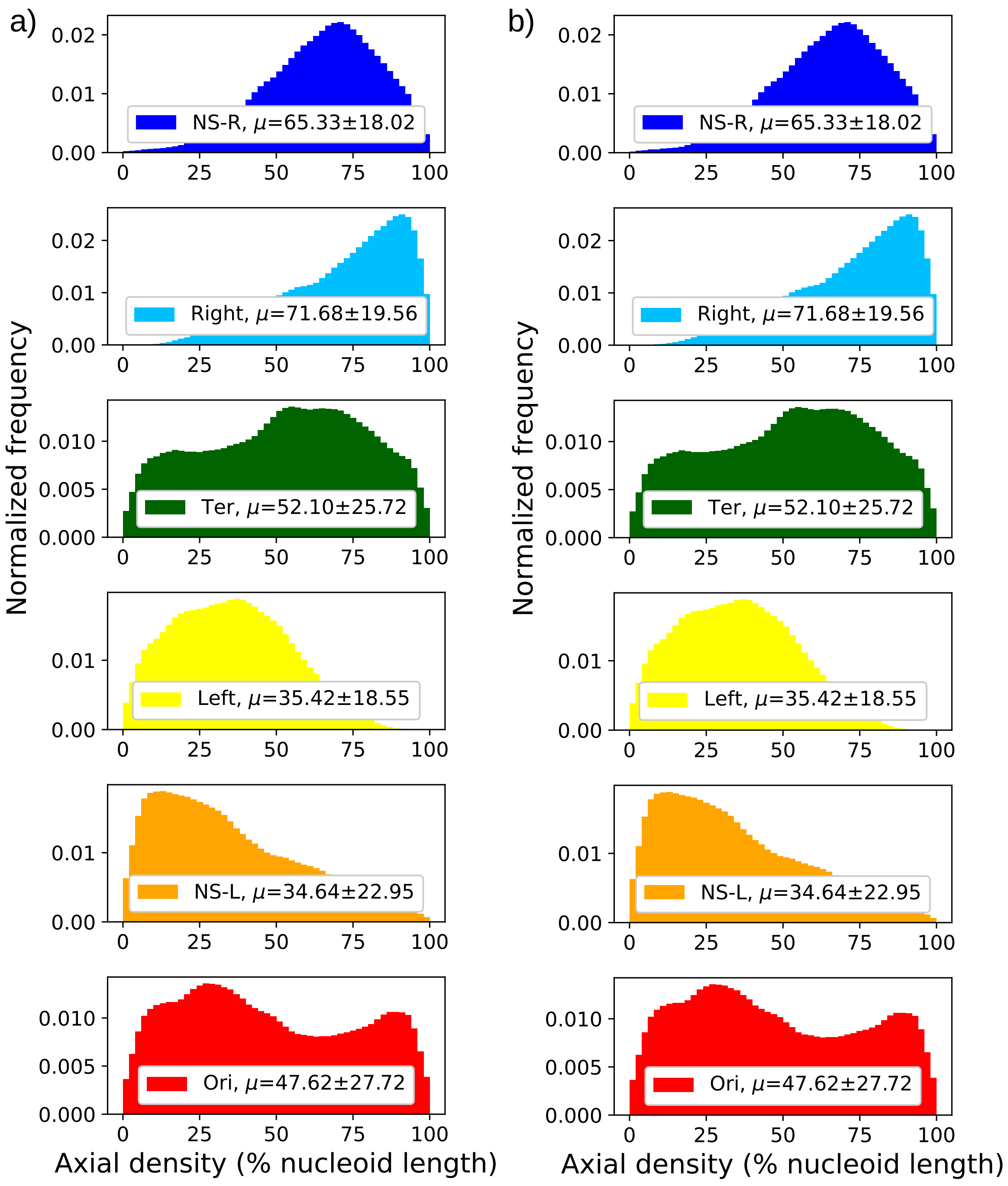


**Figure S7: a)** Axial/linear density of the macrodomains for ΔhupAB mutant *E. coli*, grown at 37°C in LB, along the long axis of the cell with respect to nucleoid length, **b)** Axial/linear density of the macrodomains for ΔFIS mutant *E. coli*, grown at 30°C in MM, along the long axis of the cell with respect to nucleoid length.

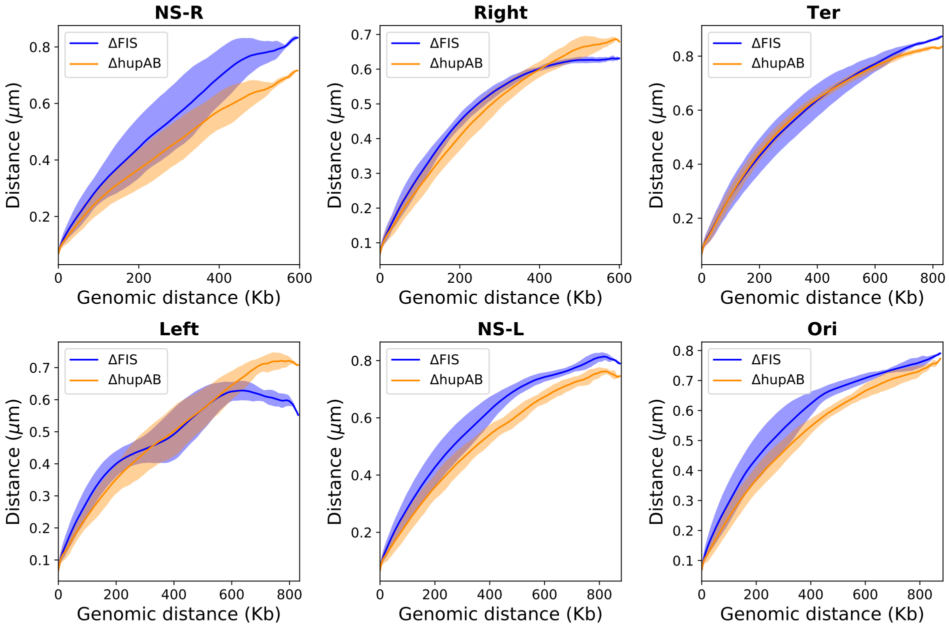


**Figure S8:** Rms end-to-end distance vs. genomic distance plot for intra-domain distances in ΔhupAB mutant (orange), grown at 37°C in LB media and ΔFIS mutant (blue) cells, grown at 30°C in MM media.

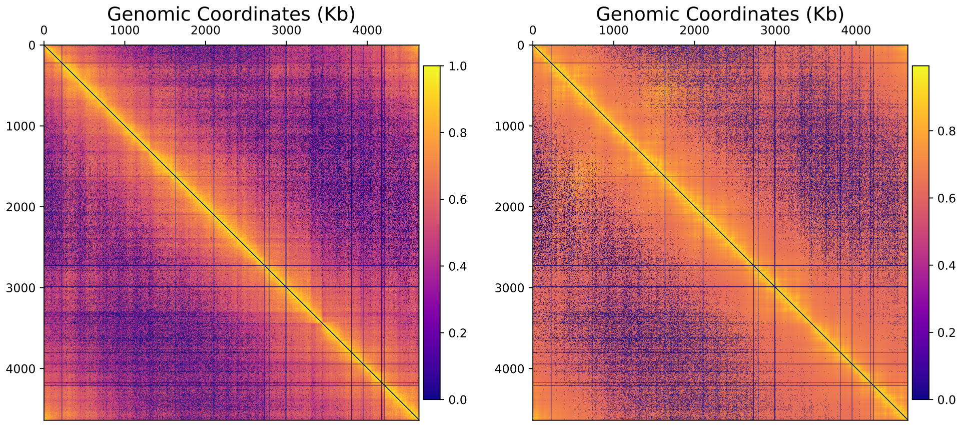


**(a)** Experimental matrix

**(b)** Simulated matrix

**Figure S9: a)** experimental contact probability matrix for ΔMatP mutant *E. coli* MG1655 in MM at $30°C$, **b)** simulated average contact probability matrix for ΔMatP mutant *E. coli* MG1655 in MM at $30°C$.

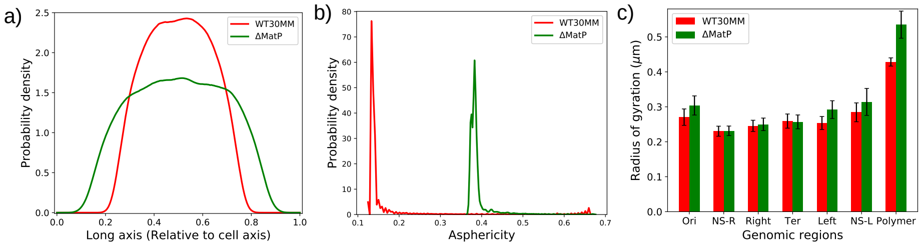


**Figure S10: a)** linear DNA probability density for WT and ΔMatP mutant cells relative to cell length along the long axis (Z- axis), **b)** distribution of asphericity of WT and ΔMatP mutant cells, **c)** radius of gyration ($R_g$) comparison between WT cells and ΔMatP mutant at same growth conditions.

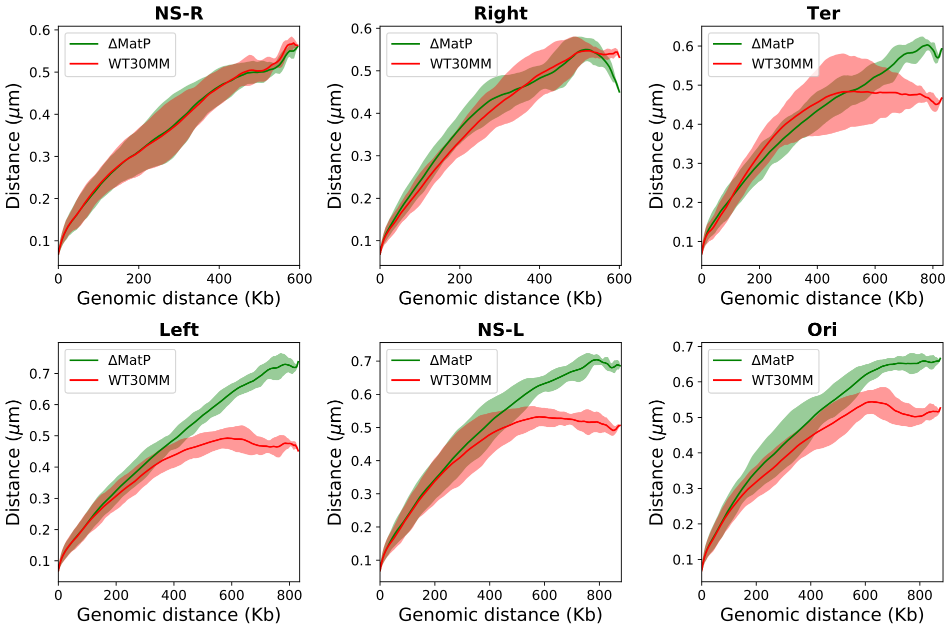


**Figure S11:** Rms end-to-end distance vs. genomic distance plot for intra-domain distances in MatP mutant (green) and WT cells at $30°C$ in MM media (red).

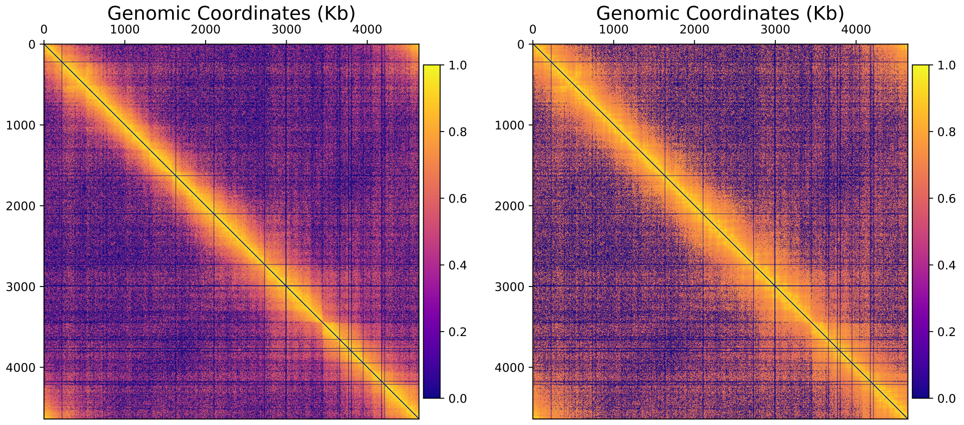


**(a)** Experimental matrix

**(b)** Simulated matrix

**Figure S12: a)** experimental contact probability matrix for ΔMukB mutant *E. coli* MG1655 in MM at $22°C$, **b)** simulated average contact probability matrix for ΔMukB mutant *E. coli* MG1655 in MM at $22°C$.

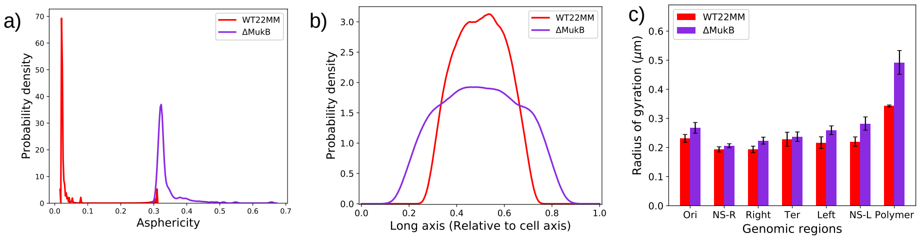


**Figure S13: a)** linear DNA probability density for WT and ΔMukB mutant cells relative to cell length along the long axis (Z- axis), **b)** distribution of asphericity of WT and ΔMukB mutant cells, **c)** radius of gyration ($R_g$) comparison between WT cells and ΔMukB mutant at same growth conditions.

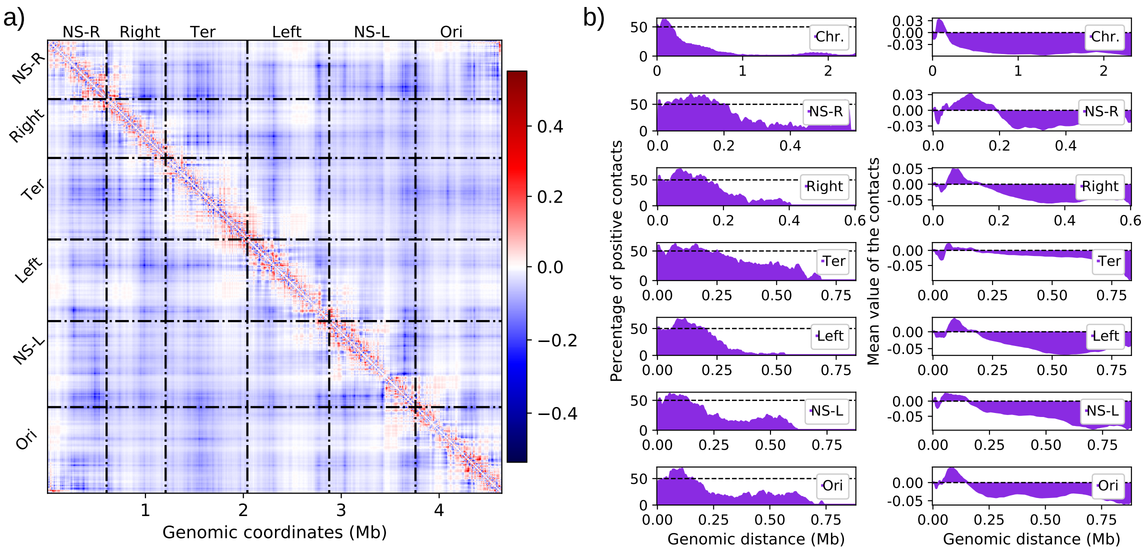


**Figure S14: a)** Difference heatmap for ΔMukB mutant (ΔMukB - WT22MM), **b) left** column: percentage of positive contacts from the difference heatmap (between ΔMukB - WT22MM) with respect to genomic distance, **right** column: average contact value from the difference heatmap (between ΔMukB - WT22MM) with respect to genomic distance.

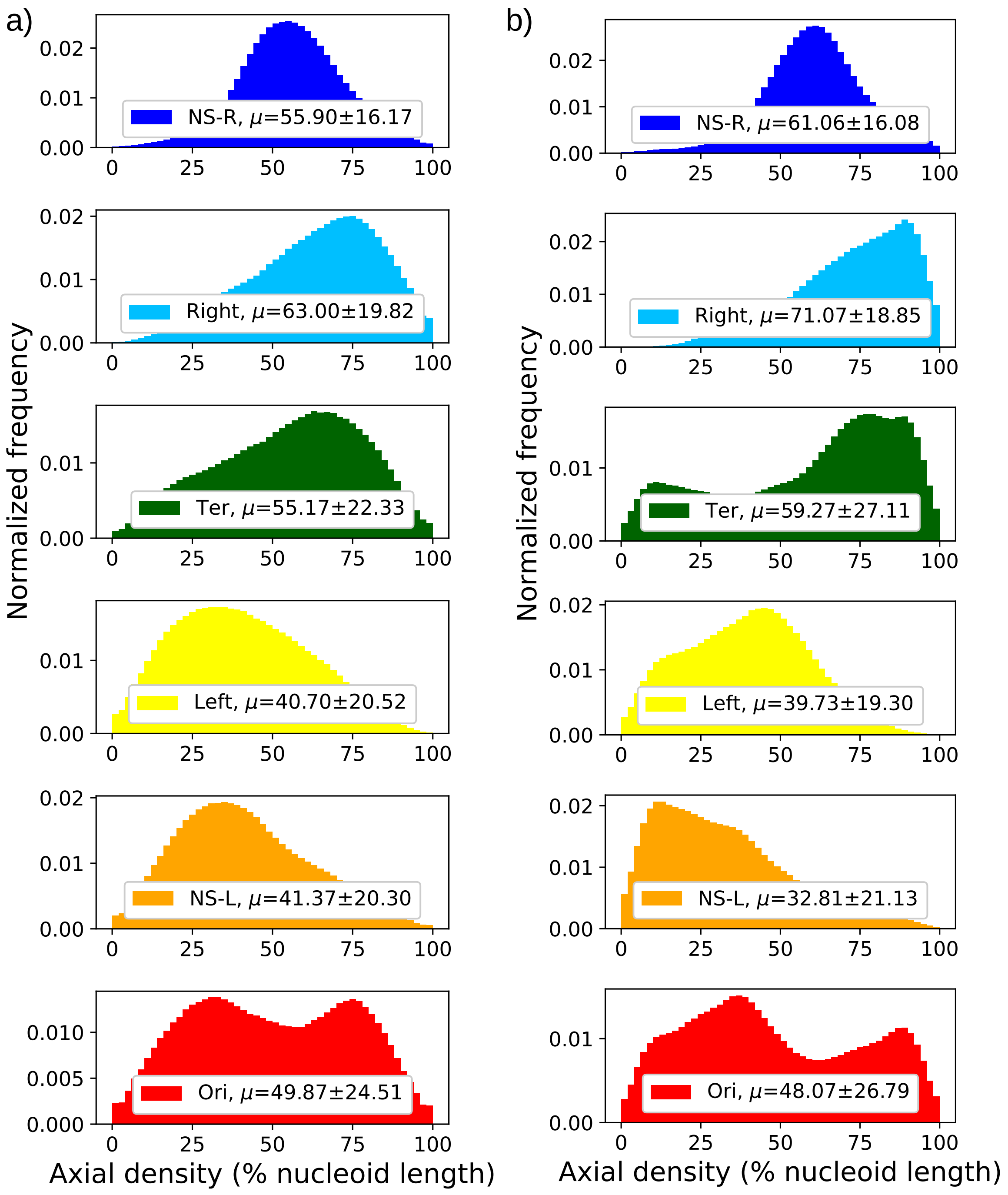


**Figure S15:** **a)** Axial/linear density of the macrodomains for WT *E. coli*, grown at 22°C in MM, along the long axis of the cell with respect to nucleoid length, **b)** Axial/linear density of the macrodomains for ΔMukB mutant *E. coli*, grown at 22°C in MM, along the long axis of the cell with respect to nucleoid length.

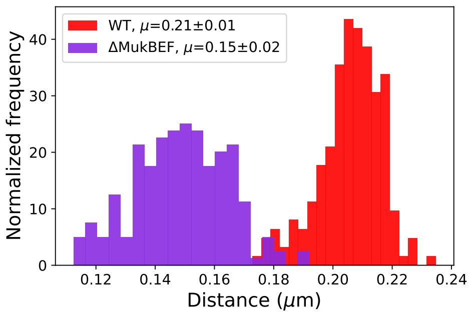


**(a)** distribution of ter2-ter4 interlocus distance

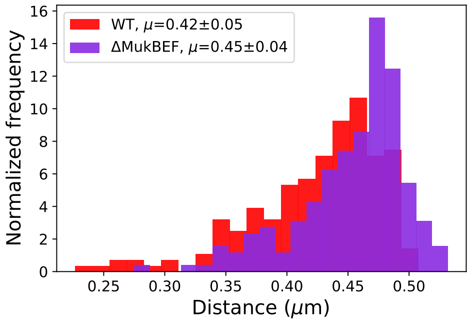


**(b)** distribution of ter2-ter6 interlocus distance

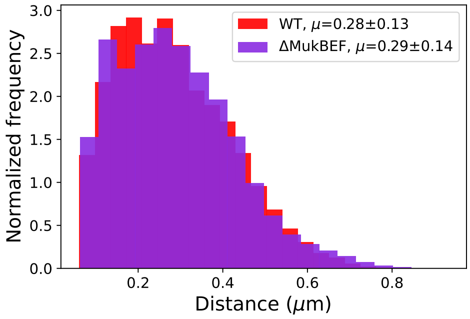


**(c)** distribution of interlocus distance between 23 matS sites pairs

**Figure S16: a)** distance between ter2 and ter4 loci which are 100kb apart in Ter region in *E. coli* chromosome for WT and ΔMukB mutant cells. **b)** distance between ter2 and ter6 loci which are ∼ 350kb apart in Ter region in *E. coli* chromosome for WT and ΔMukB mutant cells, **c)** distribution of distance between all 23 matS sites in the *E. coli* genome in WT and ΔMukB mutant cells.

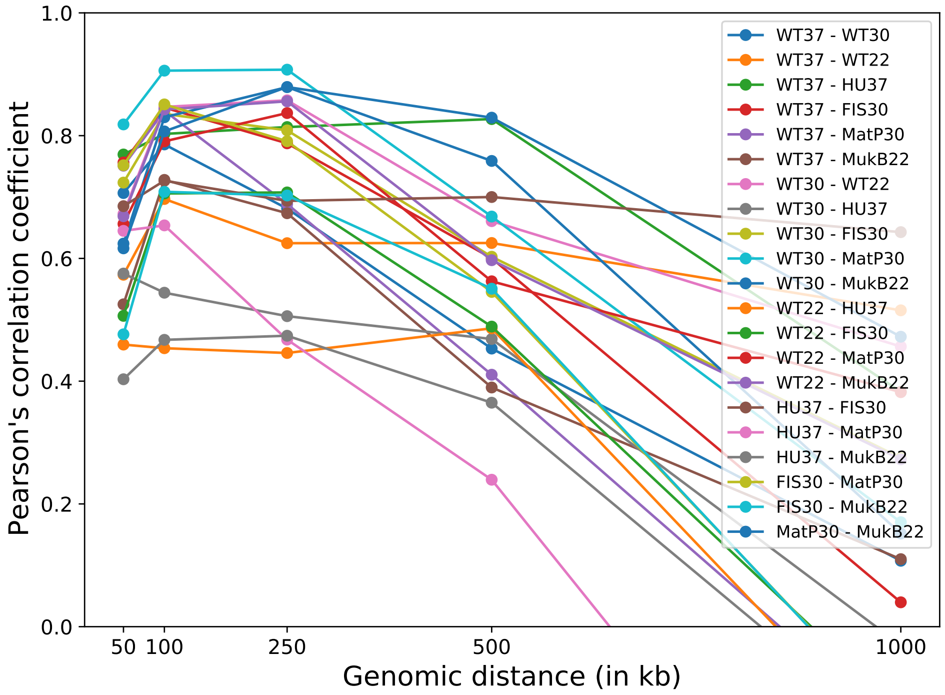


**Figure S17:** Pearson's correlation coefficients comparison between $R_g$ maps for all mutants and WT at various window sizes of 10, 20, 50, 100, 200 beads (50, 100, 250, 500, 1000kbp). The correlations are reported for pairs, where, WT37: wildtype *E. coli* at 37°C in LB, WT30: wildtype *E. coli* at 30°C in MM, HU37: ΔhupAB mutant *E. coli* at 37°C in LB, FIS30: ΔFIS mutant *E. coli* at 30°C in MM, ΔMatP30: MatP mutant *E. coli* at 30°C in MM, and ΔMukB22: MukB mutant *E. coli* at 22°C in MM. The values are given in the Table S5
